# Quantitative analysis of surface wave patterns of Min proteins

**DOI:** 10.1101/2022.04.28.489904

**Authors:** Sabrina Meindlhumer, Jacob Kerssemakers, Cees Dekker

## Abstract

The Min protein system is arguably the best-studied model systems for biological pattern formation. It exhibits pole-to-pole oscillations in *E. coli* bacteria as well as a variety of surface wave patterns in *in vitro* reconstitutions. Such Min surface wave patterns pose particular challenges to quantification as they are typically only semi-periodic and non-stationary. Here, we present a methodology for quantitatively analyzing such Min patterns, aiming for reproducibility, user-independence, and easy usage. After introducing pattern-feature definitions and image-processing concepts, we present an analysis pipeline where we use autocorrelation analysis to extract global parameters such as the average spatial wavelength and oscillation period. Subsequently, we describe a method that uses flow-field analysis to extract local properties such as the wave propagation velocity. We provide descriptions on how to practically implement these quantification tools and provide Python code that can directly be used to perform analysis of Min patterns.

## 1 INTRODUCTION

Pattern formation is a fascinating basic phenomenon that occurs across many scales, from galaxy formation all the way down to embryology and beyond. In chemistry and biology, self-organizing patterns emerge from the combination of specific intermolecular interactions and molecular transport processes.(1) Acting together, reaction and diffusion can result in inhomogeneous concentrations of molecules that constitute spatiotemporal patterns.(2) Such patterns provide useful functions as they impose directional or positional preferences on processes in cells and tissues.(3, 4, 5) Indeed, pattern formation is of vital importance for the description of a multitude of biological phenomena, ranging from bacterial and eukaryotic cell division(4), to embryonic development of multicellular organisms(3, 5), up to entire ecosystems(6).

For example, in *E. coli* bacteria, a pattern-forming mechanism acts to determine the central position of the rod-shaped cell, ensuring that the required protein-machinery is guided to the correct location to start a fully symmetric division into daughter cells. This particular pattern-forming system, formed by the Min proteins, relies on a reaction-diffusion mechanism and is widely considered to be the best-studied model system for intracellular pattern formation.(4, 7) As the moment of cell division approaches, Min proteins will periodically bind and unbind the inner membrane at the poles of the bacterium. When visualized by fluorescent labelling, the Min proteins are observed to oscillate from pole to pole with a period of approximately one minute. As a result, their temporally averaged concentration is minimized at the mid-cell location. One component of the Min system, MinC, exerts an inhibitory effect on a component of the cell division machinery. As the lowest concentration of MinC is found at mid-cell, the proteins that facilitate cell division will preferably bind there, and the cell gets divided evenly.(8)

While MinC is important for this downstream process, the pattern as such is formed by only two proteins, MinD and MinE. MinD is an ATPase that upon binding ATP can attach to the membrane, while MinE is its ATPase activator. MinE can bind to the membrane upon recruitment by membrane-bound MinD and subsequently facilitate MinD’s ATP-consuming membrane detachment.(9, 10, 11, 7) The Min proteins thus constitute a pattern-forming model system with only 2 essential components, which is appealingly simple for both theoretical and experimental studies. A wide range of experiments have been reported that explore particular features of the Min system.(12, 9, 13) A multitude of Min protein models(14, 15) have been proposed, and continue to be developed as new molecular details are discovered.(16, 17, 10, 18)

Arguably, the most iconic images of Min protein patterns present themselves in *in vitro* studies of these proteins. Such experiments typically reconstitute Min proteins on supported lipid bilayers on a glass slide, with a fraction of Min proteins carrying a fluorescent label.(9) Min proteins exhibit mesmerizing dynamic membrane patterns in such an artificial *in vitro* environment: Over a wide range of concentrations, one encounters characteristic patterns such as rotating spirals or travelling planar wave fronts, with typical wavelengths in the order of tens of micrometers.(9, 12) An example is given in Fig. 1A, showing a snapshot image for labelled MinE. Examination of this image reveals various features that are typical for Min patterns. For example, Min proteins exhibit planar waves, but these occur only locally and become less correlated on longer length scales. Min patterns thus organize in surface domains with one dominating type of pattern, such as spiral or planar wave fronts traveling in a certain direction. Boundaries between domains are recognizable by phase shifts.

**Figure 1.**
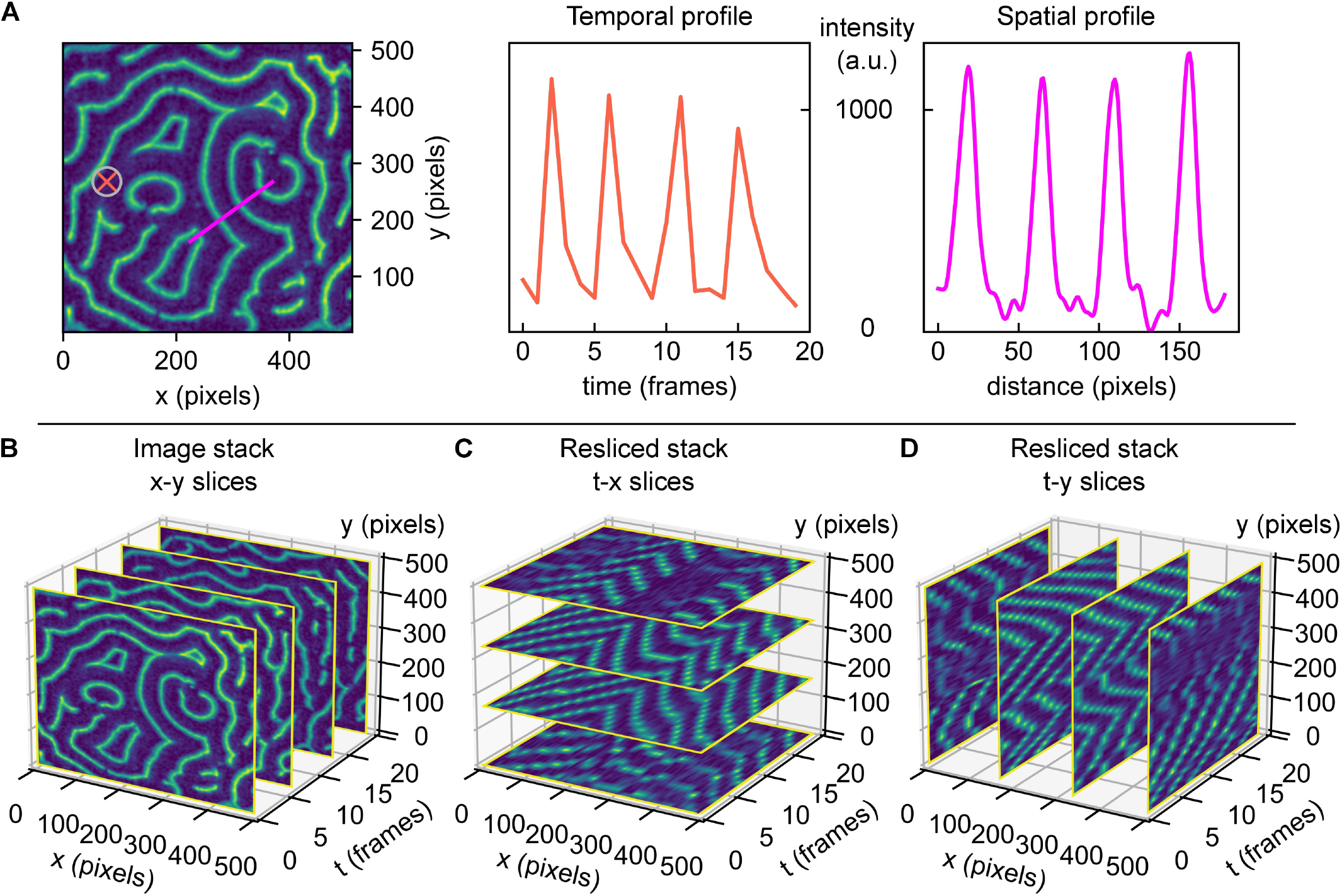
Min acquisition stacks contain spatial and temporal information on the pattern. Min protein surface patterns are dynamic in space and time. Min pattern acquisitions are 3D (*x, y, t*) matrices, containing the measured fluorescence intensity values for different coordinates. The information contained within this matrix can be accessed and visualized differently, dependent on what parameters are of interest. **A** Left: Example of a real MinE protein surface pattern. Middle: Fluorescence intensity over time at the surface position highlighted by orange cross in the image, for an image stack comprising 20 frames. Right: Fluorescence intensity trace along surface cross-section highlighted in magenta in the image. **B** 3D view of a Min protein acquisition stack with four example images (standard *x*-*y* slices) shown for demonstration. **C** 3D view of slice from A, resliced along constant *y*-directions, creating *t*-*x* slices. **D** 3D view of slice from A, resliced along constant *x*-directions, creating *t*-*y* slices.

Thorough quantitative analysis of Min patterns is nontrivial. While visual inspection is often sufficient to deduce the basic Min pattern, quantitative information is required to extract trends in the patterns’ characteristic parameters, such as the average or local wavelength, oscillation period, or propagation velocity. For perfectly periodic Min patterns with only one type of pattern with a set orientation over the entire imaged surface, analysis can easily be achieved by Fourier analysis. However, this is most often not possible due to the fact that real Min patterns are usually a patchwork of domains separated by phase-shift boundaries. The semi-periodic nature of Min patterns makes image analysis challenging, and so far there are no clear guidelines or standards on how to realise this without user bias. Extraction of quantitative parameters is typically performed by manual selection of individual surface positions,(19) line traces(20), or rectangular regions-of-interest(21, 12). Plotting of fluorescence intensity along a defined axis allows to determine the pattern’s local wavelength. Similarly, the temporal characteristics such as its oscillation period and wave propagation velocity can be measured by monitoring the intensity versus time at one spot,(19) see Fig. 1A for an illustration of these possible practices.

While these approaches are widely used, such manual selection is cumbersome, prone to user bias, and it underuses the vast amount of data in videos of Min patterns that could improve data accuracy. In the example given in Fig. 1A, the intensity profile of denoted line trace (magenta) shows approximately equidistant peaks that indicate the wavelength. However, selection of another spot or a slightly different trace that is not perpendicular to the wave fronts would have led to a deviating result. Furthermore, note how extending the trace over the region of the spiral domain would erroneously lead to the inclusion of shorter or longer wavelengths. Notably, these domain boundaries do not always remain stationary over the time of acquisition, and hence line traces would have to be carefully adapted for every single frame so as to adapt to possible reorientations of wave fronts (compare for example the domain variation over time in Fig. S1A). In view of all these limitations, we conclude that an automated user-independent analysis of large regions is preferable, as it allows to obtain solid statistics on the patterns’ characteristic features.

In this paper, we present a methodology for analyzing Min patterns that aims for thorough quantitation, easy usage, reproducibility, and user independence. We start by briefly proposing strategies for image cleaning. After that, we proceed to presenting strategies for global and local analysis of Min protein surface patterns. For extracting global parameters, we rely on calculating autocorrelation maps to examine the average periodic features of Min patterns, following other groups(16) as well as our own.(18) We treat a Min pattern acquisition stack as a threedimensional matrix containing information in space and time (*x, y, t*), and we propose strategies on how to efficiently access this information. Subsequently, we introduce an analysis pipeline which allows to quantify local properties of Min patterns, such as the wave propagation speed and direction of propagation. The approach we present here relies on the identification of individual wave crest points and their movement from one frame to the next. While other tools have been used to extract the directional preferences of Min surface wave propagation(22), our approach allows not only for obtaining large distributions of parameters, but also for accessing multiple parameters at the same time. For both global and local analysis, we offer guidance to researchers who would like to implement similar strategies for their own applications. Code is openly available and provided in Python 3(23).

## 2 IMAGE PROCESSING METHODOLOGY

Min patterns are phenomena that occur at lipid membranes. Accordingly, the most suitable forms of microscopy for *in vitro* experiments are those that acquire an imaging slab along the membrane-coated surface. Examples are total internal reflection fluorescence (TIRF) microscopy,(19, 12, 16, 24) laser scanning confocal microscopy(21, 22, 20, 25) or spinning disc confocal microscopy(18, 26). Epifluorescence microscopy(27) is generally not suitable as it leads to high background signal from the fluorescent molecules in the bulk, significantly reducing the pattern’s quality or even making it unrecognizable.

“Image cleaning” includes procedures and operations performed on microscope data that improve the overall quality of the image by making the features of interest better recognizable, while not distorting or removing essential components. Good image cleaning, where background and spurious contributions are removed, is important for successful quantitative analysis of Min protein patterns, as insufficient cleaning can lead the algorithms to fail to recognize the pattern as such. There are multiple imaging artefacts that need to be corrected. For example, local fluorescent impurities such as protein aggregates may give a static signal that is not part of the Min pattern. And depending on the microscope, illumination is typically inhomogeneous across the field-of-view. In many cases, researchers are interested in the time-dependent behavior of Min patterns, and for this, they acquire images at the same spatial positions at regular time intervals ranging from seconds to minutes. Consequently, Min proteins may bleach due to prolonged imaging.(18) Flow-cell setups with constant bulk flow may pose additional challenges in the form of objects entering and passing through the field-of-view or adhering to and spontaneously detaching from the surface.(26) We empirically found that the occurrence of imaging artefacts can vary strongly, depending on the experimental setup (specifically, the type of sample chamber and the surface preparation), protein concentration (higher concentrations may result in more aggregates), and experiment duration (the longer, the more aggregates occur).

With image cleaning, we aim to tackle all of these issues. Here, we propose a sequence of standard actions that we found to yield overall good results for a time series (several frames of the same sample position saved sequentially in an image stack) of images *I*_*movie*_ of *in vitro* Min surface patterns. Our protocol involves the following steps:

- Correct for fluorescence bleaching by normalizing each frame to its mean intensity value.
- Create an ‘illumination correction map’ *I*_*illum*_ by smoothing and averaging all movie images, and normalizing the images to their maximum intensity.
- Create a static background image *I*_*stat*_ by averaging over all the moving (surface pattern) features of all images in the stack. This background image then only contains static fluorescent features such as local specks, holes, and scratches.
- Correct each image via the following image operation: *I*_*cor*_ = (*I*_*movie*_ − *I*_*stat*_)*/I*_*illum*_.
- At this point, there may still be some artefacts left, for example, those that appeared during the acquisition time and might thus not have been included in the static background image. Therefore, remaining bright or dark artefacts can be removed manually or by thresholding.
- Images can finally be slightly smoothed to diminish effect of sharp edges and artefacts from the latter cropping.

In cases where it is required to explore Min patterns across very large areas, it may furthermore be of interest to stitch multiple fields-of-view of Min patterns into larger images.(28, 26) If the microscope’s software does not offer an automatized solution for this, stitching can be achieved by good bookkeeping and a few lines of code. When planning to stitch images, it can be helpful to choose individual field-of-views in such a way that there is a bit of an overlap (e.g. 5 % of the width) between adjacent areas, as this makes it easier to correctly reassemble the full image afterwards. Note that in general, adjoining areas may not have exactly the same median intensity. Therefore, differences in median intensity levels between individual field-of-views need to be corrected for when assembling the stitched image. This way, the resulting large image will appear more homogeneous in brightness and stitching borders will be less visible.

## 3 PATTERN ANALYSIS STRATEGIES

### 3.1 Global parameters

In many cases, researchers are interested in quantifying parameters such as the average spatial wavelength and oscillation period of a Min pattern. For this purpose, they acquire image stacks of Min patterns at a certain surface region (*x, y* in pixel or distance units) and at regular time-intervals (*t* in time units or frames). However, the existence of domains within and dynamics of patterns make manual extraction of these parameters a laborsome task, and the results of such analyses (cf. Fig. 1A) may suffer from poor statistics and user bias. Here, it is important to realize that what we are interested in are essentially global, image-averaged parameters. In this section, we describe how quantification of such global parameters can be achieved by performing autocorrelation analysis(29) for different slices along time or space coordinates (*x*-*y* frames such as in Fig. 1B, *t*-*x* slices such as in Fig. 1C or *t*-*y* slices as in Fig. 1D). We start with spatial autocorrelation, aiming to quantify the wavelength, and then continue to present how temporal autocorrelation can be used to quantify the oscillation period of a pattern.

#### 3.1.1 Global wavelength

Spatial autocorrelation analysis of an image essentially compares each pixel to other pixels around it, quantifying their (dis)similarity as a function of distance.(29) Starting from a single image frame as shown in Fig. 2A, an autocorrelation map can be calculated using routines from scientific libraries provided for most programming languages. Before performing these operations, the image should be normalized by a series of actions (subtraction of minimum intensity, division by summed-up total intensity and subtraction of mean intensity). Using standard functions supplied within most environments, a twodimensional autocorrelation map *crmx* can be calculated for an image frame *image* by the transformation

**Figure 2.**
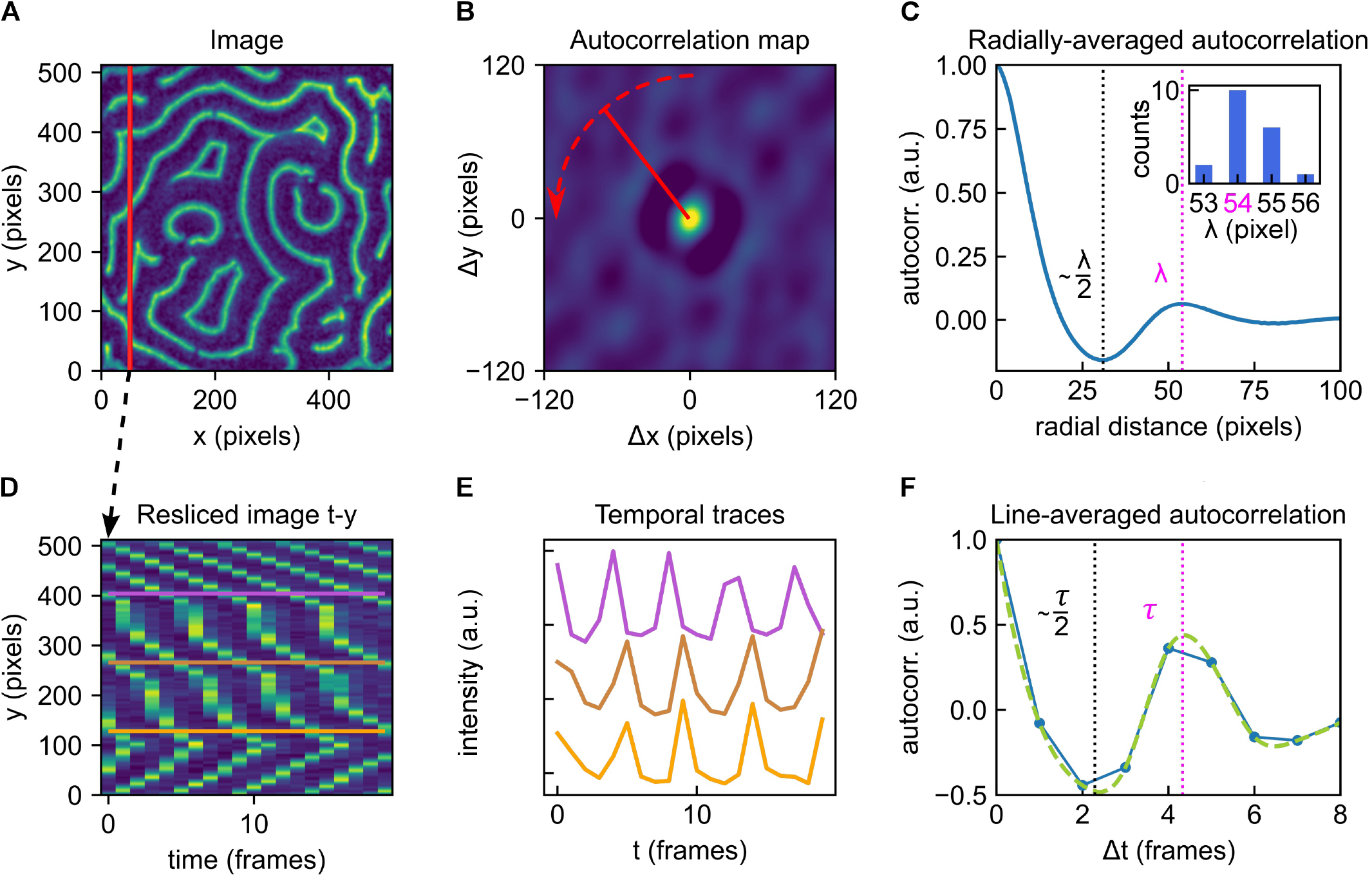
Overview of the autocorrelation analysis pipeline to obtain global parameters. **A** Example of a Min protein surface pattern, single frame. Color brightness indicates fluorescence intensity (a.u.). **B** Autocorrelation map of the frame shown in A. Color brightness indicates intensity. Red dashed line indicates radial averaging. **C** Averaged radial profile for the autocorrelation map shown in B. The first maximum after the central peak at distance zero is a measure of the the patterns global characteristic wavelength. The identified wavelength λ is highlighted by the magenta dashed line. The first valley is positioned at approximately λ/2 (black dashed line). Inset: the analysis presented in A-C can be performed for all individual frames within an image stack. Here, the collected results for 20 consecutive frames (image stack also represented in Fig. 1B) are shown in a histogram. A λ of 54 pixels is deduced. **D** Resliced image for fixed *x*; same as shown in Fig. 1D. **E** Selected temporal traces for the *y*-positions indicated in D. All point-traces shown in the kymograph in D will be included in the autocorrelation trace shown in F. **F** Line-averaged autocorrelation trace from point-traces (such as shown in E) in blue, cubic spline fit used for peak-detection in green. The first maximum after the central peak at Δ*t* = 0 measures the average characteristic oscillation period of the pattern. The identified global oscillation of *τ* = 4 frames is highlighted by the magenta dashed line. The first valley is positioned approximately at *τ/*2 (black dashed line). The analysis pipeline illustrated in D-F can be performed for multiple slices (fixed *x* or *y*) of the image stack.

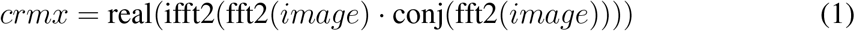

with *fft2* calculating a two-dimensional discrete Fourier Transformation, *ifft2* calculating its inverse and *real* and *conj* returning the real part and complex conjugate of the input, respectively. The four quadrants of the autocorrelation map can be re-arranged so as to place the position (Δ*x* = 0, Δ*y* = 0) in the center of the image, rather than having four partial peaks in the corners.

Close examination of the autocorrelation map in Fig. 2B reveals a general speckled pattern as well as a ring-shaped feature that is present around the center, which is characterized by a peak in intensity. As this central peak corresponds to a distance of zero, its high intensity is a consequence of the self-correlation of each pixel with itself. Moving radially outward from the center, one observes a decrease in intensity (negative autocorrelation) followed by a peak (positive autocorrelation). An easy way to extract the pattern’s dominant wavelength is to perform an angular averaging of the profile around the central peak, where the radial line profiles are averaged over all angles, as illustrated by the red arrow in panel Fig. 2B. This leads to a profile as shown in Fig. 2C. The first minimum corresponds to the dark value from the autocorrelation map. Here, the first peak after the central peak identifies the pattern’s global wavelength λ (magenta dashed line). The valley in fact corresponds to ∼ λ/2 (black dashed line). Note that this value is not exactly half the wavelength, since Min profiles are typically somewhat asymmetrical (cf. Fig. 1A).

If, as often is the case, Min patterns display steady-state dynamics, then multiple time frames can be used to provide better statistics in estimating the global wavelength. Min patterns are routinely acquired at constant time intervals, which provides a series of sequential image frames, as represented in Fig. 1B. Calculating and analysing autocorrelation maps for several frames within such an image stack allows to collect a distribution of wavelengths, such as the histogram shown in the inset in Fig. 2C, summarising results for 20 consecutive frames. Indeed, for dynamic patterns, we recommend performing this analysis strategy for multiple (e.g. 10) frames and averaging the results to obtain a reliable value for the pattern’s spatial wavelength. As Min patterns are typically dynamic, it is also possible that their wavelength changes over time, e.g., due to changes in external parameters. The spatial autocorrelation analysis presented here is however equally applicable to static patterns.

Identifying the peaks shown in Fig. 2C (and 2F, see following section) is trivial to the human eye, but not necessarily straightforward to automate. For our data, we achieved this via a simple algorithm that identifies the second local maximum of the trace. In doing so, we implicitly assume that the trace is fine enough to achieve sufficient accuracy and upsampling is not required. While a curve such as shown in Fig. 2C looks smooth to the eye, the original image data had a finite resolution, and accordingly, this curve contains small rags which an algorithm may erroneously detect as a local maximum. In many cases, this issue can be solved by smoothing the curve slightly before subjecting it to peak analysis. This can be achieved by applying a smoothing kernel stretching over a few pixels, for example a small fraction of the image dimension.

To avoid erroneous results, it is recommendable to plot at least one example trace (for one frame) and inspect whether the detected (averaged) wavelength position correctly co-localizes with the valley position. If no clear valley can be identified (characterized by a small or even vanishing intensity difference between the local minimum and maximum), the pattern may lack a recognizable periodicity, which could be an intrinsic property of the given pattern. Alternatively, this issue can also be encountered if insufficient image cleaning has been applied to the stack before analysis and the pattern is superimposed with too many artefacts, resulting in low autocorrelation. However, we find that this method is generally quite robust towards noise, as shown in Fig. S2B.

Further, the ratio between image size and spatial wavelength is of importance. In Fig. S3, a large-scale image stack is sequentially cropped to smaller sizes and global autocorrelation analysis as presented here is performed on randomly selected areas. Based on these results, we find that the standard deviation increases as the image sizes gets smaller, and estimate that the image dimension should be large enough to contain at least 5-10 full wavelengths for the presented algorithm to provide reliable results.

#### 3.1.2 Global oscillation period

Another global property of interest is the average oscillation period. This parameter describes how at any given point within the imaged region, the membrane protein density (which is proportional to the fluorescence intensity) can be expected to change over time. We quantify the global oscillation period of a Min pattern following a strategy that closely resembles the one presented for obtaining the global wavelength in the preceding section.

To obtain quantitative information on the temporal dynamics of Min patterns, Min patterns are typically acquired as time series. A time series such as the one shown in Fig. 1B is essentially a threedimensional (*x, y, t*) matrix, containing information on fluorescence intensity as a function of space and time. Notably, for temporal analysis to be reliable, image data has to be acquired at a rate above the Nyquist rate. Min patterns are characterized by point-wise temporal periodicity (compare the time evolution at a random sample position in Fig. 1A). Hence, when acquiring a time series, intervals have to be chosen short enough to ensure the acquisition of enough data to adequately depict the pattern’s temporal dynamics. If consecutive acquisition times in a series are chosen very far apart, the pattern may even appear to travel towards the opposite direction. According to Nyquist theorem, for patterns of an oscillation period *τ*, consecutive images in a series have to be acquired at intervals spaced no longer than *τ/*2 apart. In principle, the shorter these intervals are, the better the temporal dynamics can be characterized. Min protein patterns in lipid bilayer reconstitutions will typically exhibit wave propagation velocities of several 100 nm s^*−*1^(12) and wavelengths around 50 µm(9). This leads to an expected oscillation period in the order of minutes. Acquiring sufficient images to fulfill Nyquist theorem should therefore not be an issue in most microscope setups. In practice, finding the optimal imaging rate is a trade-off between achieving a high temporal resolution and minimizing fluorescence bleaching.

Upon analysing a given Min pattern stack with respect to its temporal characteristics, we found that reslicing the stack along fixed spatial coordinates simplifies data processing and makes it more straightforward to access the temporal information contained within the stack. Fig. 2D shows a (*t, y*) slice through our example image stack at fixed pixel position *x*. These *t*-*x* or *t*-*y* slices are collections of kymographs for all points along the cross-section at the fixed spatial coordinate. In Fig. 2E, we show example traces of these kymographs along selected *y* positions, which then correspond to individual surface locations (*x,y*) on the field-of-view shown in Fig. 2A. Analogous to the strategy presented for spatial global analysis, Eq. (1) can be used to obtain an autocorrelation map for this slice (cf. Supplementary Appendix *DEMO_MinDE_global_analysis*). To extract the pattern’s global oscillation period, we consider the autocorrelation map’s trace at Δ*y* = 0 for Δ*t* ≥ 0, plotted in Fig. 2F. Including the Δ*t* ≤ 0 trace (going back in time) of the curve would not yield additional resolution, as it contains the same information as the positive trace (going back in time). This profile then immediately provides the averaged temporal autocorrelation trace for all points along one fixed spatial coordinate (as indicated for fixed *x* by the vertical line in Fig. 2A). If the line-averaged autocorrelation trace does not provide sufficiently high resolution to identify a minimum indicative of the oscillation period, imaging over a longer time and/or at shorter intervals could be necessary.

The first maximum of this curve beyond the central peak again identifies the predominant global oscillation period *τ* (magenta dashed line), while the first minimum is located at approximately *τ/*2 (black dashed line). Statistics on *τ* can be collected by performing the analysis pipeline illustrated in Fig. 2D-F for multiple cross-sections at fixed *y* or *x*, that is, for multiple resliced frames as shown in Fig. 1C and 1D. Compared to the traces obtained from spatial autocorrelation, which are calculated via radial averaging, these temporal autocorrelation traces may appear rather coarse. A more advanced peak detection can be obtained with cubic-spline fitting and subsequent extreme point analysis. Further, the presented method for temporal autocorrelation analysis is less robust towards noise than spatial autocorrelation, as shown in Fig. S2C. Adequate image preprocessing and cleaning is again of high importance. Image smoothing was found to slightly increase the detected oscillation period, albeit not above the error margins obtained for unsmoothed images, as shown in Fig. S4.

Note furthermore that the propagation speed of a pattern may depend on the concentrations of all components, and may change over time. Therefore, for very long time series, it can make sense to split the stack up into multiple stacks of shorter total duration and perform analysis for subsequent sub-stacks (compare Fig. S8). However, we estimate that an image stack needs to span the duration of a minimum length of 3 oscillation periods for the presented method to yield reliable results (cf. Fig. S4).

### 3.2 Local parameters

In this section, we present an analysis strategy that aims to quantify local parameters of Min patterns. These parameters include the local wave propagation velocity or characteristic local distances such as the distance between MinD and MinE wave crests. As we will describe in more detail, local parameters can be quantified by implementing two steps: (i) identification of wave crests for each frame within an image stack, and (ii) comparison of intensities at and around the identified crest points from one frame to the next.

#### 3.2.1 Local wave propagation velocity

The local wave propagation velocity is determined as the shift of individual crest points from one frame to the next. To calculate the local velocity, we first need to identify these crest points, and then determine their positional difference. We achieve this by applying optical flow analysis. Optical flow is defined by how brightness patterns move from one image to another(30, 31). For our Min patterns, we estimate an optical flow vector field using Horn-Schunck algorithm(30, 32). This algorithm works on a pair of sequential image frames (see Fig. 3A for example data, recorded for MinE) and returns a vector field, describing the estimated optical flow for each pixel from the first to the second frame. Fig. 3B shows the magnitude (indicated by image brightness) of the optical flow vectors obtained for our example image pair from Fig. 3A. Note that the collected optical flow vectors do not necessarily represent the local wave propagation velocities. The relation between the two measures will depend on the wave shape, making it hard to generalize.

**Figure 3.**
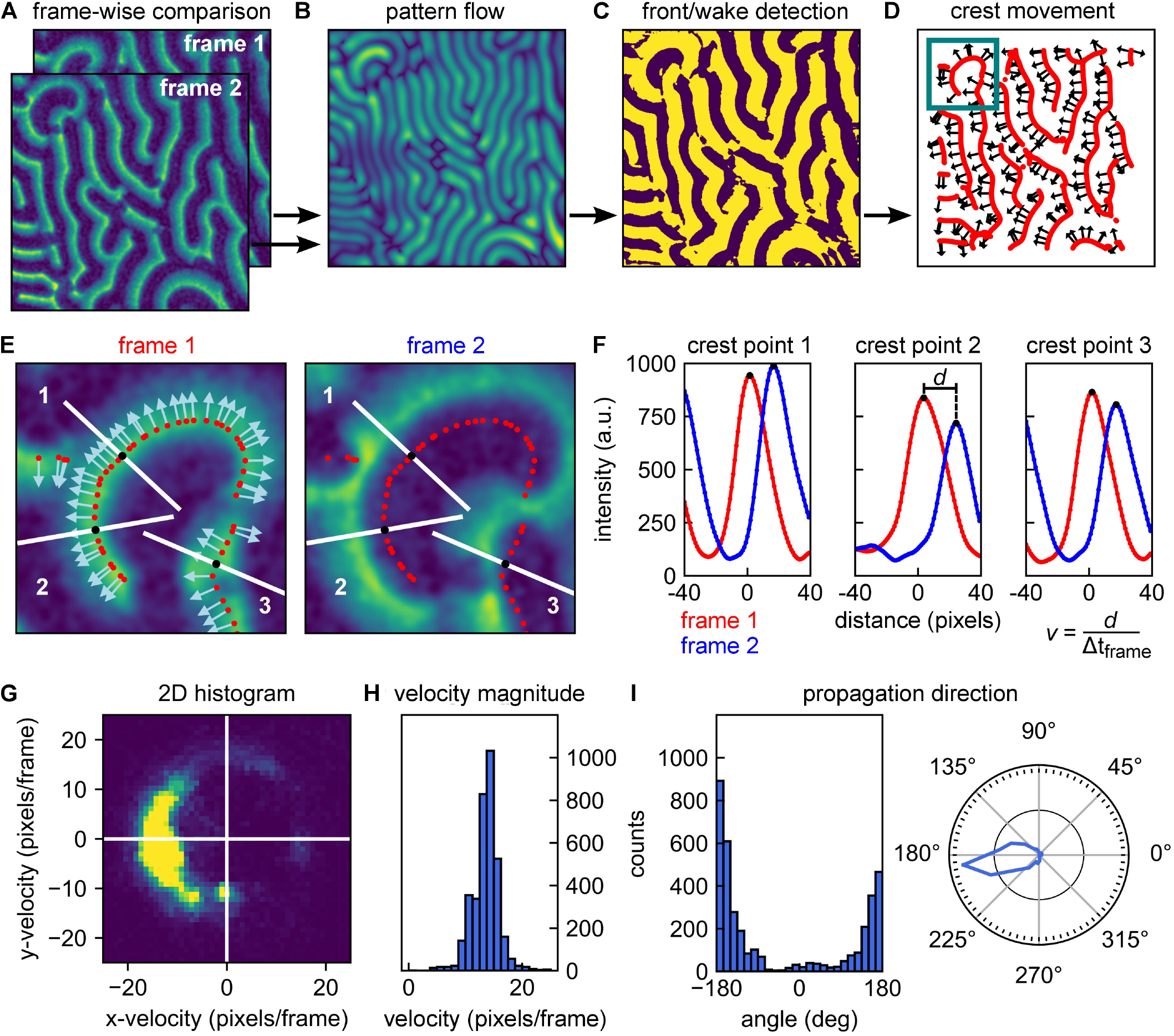
Overview of analysis pipeline for determining local wave propagation velocities. **A** Local analysis relies on sequential pairwise comparison of frames within an image stack. Example frames are here referred to as frame 1 and frame 2. **B** the Horn-Schunck algorithms is used to compute optical flow of the image pair (from frame 1 to frame 2) These return one vector per pixel, indicating the shift. Brightness indicates the vector magnitude. **C** Phase image of frame 1. The phase is determined by whether the wave intensity goes up (“front” of the wave, bright color) or down (“wake” of the wave, dark color) following the direction of the vector obtained from the optical flow analysis. **D** Wave crests can be identified by the transitions from front to wake as denoted in panel C. Red lines show the identified wave crest locations. Selected arrows indicate the direction of movement of the crests as determined from the optical flow. **E** Left: Zoom-in of frame 1 at the location highlighted in panel D. Wave crest points are indicated by red dots; direction of movement as identified by optical flow analysis are indicated by arrows. Right: Zoom-in of frame 2 at the same location. Crest points of frame 1 are indicated by red dots. In both images, 3 crest point positions are highlighted by white line traces parallel to the direction of propagation at these points (i.e., normal to the wave fronts). **F** Intensities versus distance along the white lines in panel E. Starting from the crests identified for frame 1 (red dots), intensities are sampled at discrete sub-pixel positions perpendicular to the wave front. Smoothed intensity traces are shown in red for frame 1 and in blue for frame 2. The translocation *d* of each crest position is then determined from the peak shift, allowing to calculate a velocity vector with magnitude *v*. Wave crest velocities can be calculated for each crest and each sequential pair of frames in an image stack. **G** Velocity distribution of a pattern can be visualized as a 2D histogram. **H** Velocity histogram that provides the average velocity magnitude *v* as well as the distribution of velocities within a given pattern. **I** Histogram of angles of the wave propagation vector (left) and an angular diagram denoting the occurrence of various pattern velocities (right). From the list of velocity vectors, we have immediate access to the propagation directions present within a pattern, the occurrence of which can be visualized in a histogram (left). The information can be represented as a polar histogram (right).

As a first step to identifying the positions of the wave crests within the first frame, we use the information obtained from optical flow analysis to create a phase image. This phase image is essentially a binary map over the whole image, which indicates whether at any given point we are at the “front” (‘ahead of the wave’) or “wake” region of the wave (‘behind the wave’). These regions are characterized by a decrease or increase in fluorescence intensity, respectively, along the direction of movement. If we consider any specific surface location and its corresponding vector obtained from optical flow, we can follow the direction of that vector over a short distance (say 3 pixels) within the same frame, and determine whether the fluorescence intensity is decreasing or increasing. The outcome of this procedure for our example frames is shown in Fig. 3C. Using this phase image, we can identify those positions in the image that make up the wave crests – which form the backbone structure of the pattern. Wave crests are identified by the positions at which a transition from the front to a wake region occurs. Fig. 3D shows the wave crests identified for the first frame in red. Subsequently, we need to determine the direction of wave propagation for these individual crest points. While we have a set of vectors available for distinct (*x, y*) positions from optical flow analysis, their directions will in general not be exactly normal to the wave crests due to noise. Therefore, we calculate vectors which are perpendicular to the lines of wave crests, and determine their correct directionality (pointing towards the front of the wave) using the information already present in the phase image. In Fig. 3D the direction of propagation is indicated by vectors at distinct crest points throughout the image for illustration. Following the procedure outlined here and in Fig. 3A-D, we thus effectively obtained (i) a list of positions, identifying the wave crest point positions, and (ii) a list of vectors, describing their direction of movement from the current to the next frame.

The next step is to quantify how far these crest points moved during the transition from the current to the next frame. Upon quantifying this positional shift, the velocity magnitude is given by this distance divided by the time that passed between the acquisition of subsequent frames. The procedure is illustrated in Fig. 3E-F for three example crest points. For each crest point, we considered the fluorescence intensity traces along the direction of movement at the given position (i.e., perpendicular to the wave front). Starting from the crest position, we plotted this trace up to a distance corresponding to roughly half the (global) wavelength of the pattern. Choosing this distance should ensure that the next frame’s peak is still within the sampling range (provided that the time resolution of the acquisition stack is sufficiently high, cf. section 3.1.2). In Fig. 3E, the example traces along which intensities are plotted are highlighted in white. For sampling positions along this trace which are located off the pixel grid, intensities can be estimated by interpolation. This yields an intensity trace at sub-pixel resolution. This intensity trace will have a maximum located very close to the center of the originally determined crest position, with a small offset that can be attributed to the additional smoothing step performed before optical flow analysis, and is generally not significant (cf. Fig. S5 and adjoined table).

Next, we performed exactly the same procedure at the same positions on the next frame. As shown in Fig. 3E and Fig. 3F, these positions are now no longer centered around the maximum of the intensity trace, but offset from them by a certain distance. This distance is the positional shift that we need to obtain for quantification of the local wave propagation velocity. Performing peak fitting allows to identify the two traces’ peak values and measure the shift *d* in crest peak position at sub-pixel resolution.

Performing this analysis for all identified crest points along all sequential pairs of frames in an stack eventually yielded a list of velocity vectors with components (*v*_*x*_,*v*_*y*_) at frame *n* for a wave crest at position (*x,y*). Image smoothing is an important technical aspect of the local analysis presented here. We first perform a light image smoothing (with a limited initial smoothing kernel) to de-noise the image and to simplify peak detection such as shown in Fig. 3F. For optical flow analysis (Fig. 3B), we perform an additional smoothing step with a more extensive secondary smoothing kernel. The effect of different initial smoothing kernels is illustrated in Fig. S6, the effect of increasing image noise in Fig. S2D and E. Further, applying Horn-Schunck algorithm(30) requires setting a few parameters, particularly the regularization constant and the number of iterations. Here, we determined their values by trial-and-error from evaluating simulated as well as real Min protein pattern data and kept both constant from then on.

Depending on what kind of information is of interest, the obtained distribution of velocity vectors can be represented in different ways. A representation that combines information on the magnitude and directional preference is a 2D histogram such as the one shown in Fig. 3G for an example image stack of MinE data. If the magnitude of the wave propagation velocity is most important, the velocity magnitudes *v* can be plotted in a histogram to visualize their distribution and peak value, as shown in Fig. 3H. In other cases, the directional information may be of prime interest, for example, when studying the influence of an external bulk flow (26) or physical environmental parameters on the pattern.(22) For each vector component, an angle *α* can be calculated that corresponds to the wave crest point’s direction of movement with respect to the direction of applied flow. The occurrence of different directions can then be plotted in a regular or polar histogram, as shown in Fig. 3I. As shown Fig. S7, the method can be applied to a wide range of image sizes. Decreasing the area on which analysis is performed generally leads to smaller standard deviation in velocity magnitude distribution, which is linked to a loss in directional information.

#### 3.2.2 Local MinD/MinE crest shift

When studying Min protein patterns, researchers routinely acquire data in two channels, for fluorescently-labelled MinD and MinE, respectively. Both these proteins exhibit a peak in their surface density, but these two peaks do not coincide in space and time (an example is given in Fig. 4A).

**Figure 4.**
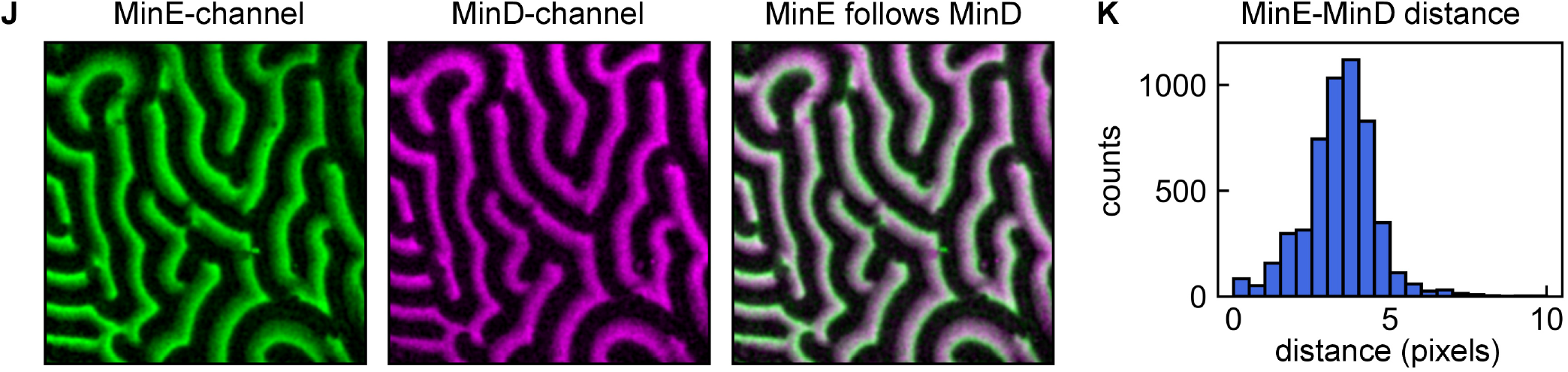
Using the local analysis pipeline to determine local MinE-MinD crest positions shifts. **A** MinD (magenta) and MinE (green) patterns. The pipeline for local analysis shown in panels A-F can be used to quantify other properties of the pattern, such as the MinE-MinD crest point distance in a simultaneous dual-channel acquisition. Here, crest point detection was performed for each frame of the MinD stack. Subsequently, the intensity traces at these positions were compared to those of the corresponding MinE frame. **B** Distribution of MinE-MinD crest point distances.

To determine these differences, the analysis pipeline presented in section 3.2.1 for obtaining wave propagation velocity between sequential frames can be slightly adapted for measuring the difference in crest position between MinD and MinE waves. This can be achieved by first performing crest detection (analysis pipeline shown in Fig. 3A to D) for one stack, e.g. the MinD stack. The crest points attributed to the MinD stack can then be compared to the corresponding frame of the MinE stack, analogous to the comparison shown in Fig. 3E. The collected peak shifts *d* can be visualized in a histogram, such as shown in Fig. 4B for our example data.

## 4 DISCUSSION

In this paper, we proposed protocols that can be applied to analyze data on *E. coli* Min protein surface patterns with respect to quantitative parameters such as the spatial wavelength, oscillation period, or wave crest propagation velocities. Our primary goal was to develop user-independent and largely automatized strategies for pattern analysis in order to (i) improve comparability of results obtained from different studies and research groups, and (ii) provide tools for quantification of experimental data for comparing results to theoretical studies. We presented global and local analysis methods which both can be valuable depending on what kind of pattern information is of interest. All of the routines that we suggested and described can be further modified to better fit the requirements posed by a particular research question – thus providing a framework that others can build upon.

The methodology we describe can be combined with many of the aspects and practices of Min pattern analysis which we did not cover here. For example, in studies investigating absolute or relative local concentrations or concentration gradients of MinD and MinE on the surface, fluorescence calibration can be important. Calibration allows to relate fluorescence intensity to the local membrane protein density.(19, 21, 12, 16) However, in many cases the patterns as such – their shapes, domains, wavelength, or dynamics – can be studied without this step, using the acquired fluorescence intensity data in arbitrary units, and this is what we focused on in this paper.

The global analysis that was presented here provides parameters averaged over entire image frames (spatial analysis) or slices along fixed spatial coordinates (temporal analysis). We found that for well-cleaned images, spatial autocorrelation robustly quantifies the main dominant wavelength within an image frame, as identified by the distance at which maximum self-correlation is present. This yielded results that are very comparable to those obtained by manual quantification, albeit with higher statistical reliability (see Fig. S1 for an example comparison of manual and automatic analysis). For temporal autocorrelation, we recommend re-orienting the image stack into slices of fixed *x*- or *y*-coordinates and calculating autocorrelation-maps along sets of kymographs, as described. We found this to be an efficient way to access large parts of the information provided within the image stack. Pixelwise analysis of intensity traces at distinct surface location, autocorrelation thereof and subsequent averaging is a more straightforward approach. However, we found that taking this detour via reslicing as presented in this paper reduces computational time by about two orders of magnitude, and hence, allows for easier processing of large amounts of data (cf. SI).

In many cases, parameters obtained from spatial and temporal analysis can be averaged for several frames/slices. However, whether this is justified depends on the time evolution of the pattern over the course of the acquisition. Fig. S8 shows the temporal autocorrelation results for an example image stack which shows a decline in oscillation period over time. This is visible from the spreading of peaks in the resliced image (cf. Fig. S8A) as well as in the variation in the shape of the Δ*x* = 0 traces (cf. Fig. S8B). In such a case, averaging is only justified over the part of the stack that shows a constant temporal periodicity. If required, the routines can be expanded to include peak analysis of single traces in the kymograph set and automatically determine up to which point averaging is permitted.

Local analysis, in contrast to global analysis, directly provides large-number distributions of parameters even when analysing only two consecutive images. Our approach to local analysis relies on the identification of wave crests and quantification of the positional shift thereof from one frame to the next. The first step can in principle be achieved by various strategies. For example, one could implement an algorithm that relies on thresholds of fluorescence intensity to identify peaks. Here, we presented a strategy that relies on established algorithms for optical flow analysis, as it offers certain advantages, such as direct access to the vector fields. We have already successfully used this local analysis methodology to examine and quantify the response of Min protein patterns to hydrodynamic bulk flow.(26) Numerous other applications can be imagined, in which a surface pattern is either exposed to an external stimulus and responds to it, or in which the time evolution of its characteristic properties is relevant. Using the local analysis pipeline presented here (or modifications thereof) allows to systematically quantify properties of interest with proper statistics rather than relying on qualitative description alone.

As our local analysis routine essentially offers a way to acquire a high number of intensity traces that are locally perpendicular to the wave fronts, it opens up possibilities for further in-depth analysis. Our brief illustration of a MinD-MinE crest distance detection is only one example of a parameter that can be extracted using an optical-flow-based routine. For example, it is imaginable to modify our local analysis pipeline such that it facilitates fast visualization of local fluorescence intensity profiles for both MinD and MinE waves. Upon fluorescence calibration, the fluorescence intensity can be converted to membrane protein density, and hence the relative peak heights can be connected to the protein ratio at the surface. Further, information obtained from local and global analysis can be combined, and local velocity analysis can provide an alternative to global methods if their requirements are not met. For example, while we found that a certain minimum number of entire oscillations is required for global temporal analysis to yield results (cf. Fig. S4), local analysis can already be performed on only two consecutive frames, and can be combined with global spatial analysis to serve as an alternative route to determining the predominant oscillation period.

Finally, we note that the general strategies presented within this paper are not restricted to application to the analysis of *E. coli* Min protein surface patterns. Min protein patterns have some properties that are very specific to them – such as their quasi-periodic nature, dynamic oscillatory behavior, and their organization within domains – and our routines are designed for these specific properties. However, the routines presented here may very well be modifiable and applicable to quantify other types of semi-periodic surface patterns, biological or not, for example to the fascinating reaction-diffusion patterns of chemical systems like the BZ-AOT system(33). We hope that our analysis toolbox will help to further disentangle the intriguing mechanisms that underlie pattern formation.

## CONFLICT OF INTEREST STATEMENT

The authors declare that the research was conducted in the absence of any commercial or financial relationships that could be construed as a potential conflict of interest.

## AUTHOR CONTRIBUTIONS

S.M. contributed to the provided image analysis tools and their representation, created the figures and co-wrote the paper. J.K. identified suitable image analysis methods, developed the majority of the provided tools and co-wrote the paper. C.D. co-developed the underlying methodology and co-wrote the paper.

## FUNDING

We acknowledge the BaSyC consortium for funding.

## ACKNOWLEDGMENTS

We thank Grzegorz Pawlik for providing example Min data, Julian Hofer for support with Python coding, and Alberto Blanch Jover, Angel Goutou, Ashmiani van den Berg, Eli van der Sluis, Erwin Frey, Fridtjof Brauns, Jaco van der Torre, Jernej Rudi Finžgar and Nicola De Franceschi for helpful discussions.

## CODE AND DATA AVAILABILITY STATEMENT

The code created for this study as well as example data are openly available and can be found on GitHub at https://github.com/M-Sabrina/MinDE_analysis_2022 and Zenodo at https://zenodo.org/record/6724666 (second release). Please refer to Supplementary Material for further information.

## Notes

https://zenodo.org/record/6724666

### Competing Interest Statement

The authors have declared no competing interest.

### Summary of Updates

In this revision, we aimed to further clarify several intermediate steps in the analysis pipelines (both global and local). New supplementary figures explore the accuracy and limits of our methods.

## REFERENCES

1. Turing A. The chemical basis of morphogenesis. Philosophical Transactions of the Royal Society of London. Series B, Biological Sciences 237 (1952) 37–72. doi:10.1098/rstb.1952.0012.

2. Bois JS, Jülicher F, Grill SW. Pattern Formation in Active Fluids. Physical Review Letters 106 (2011) 028103. doi:10.1103/PhysRevLett.106.028103.

3. Kondo S, Miura T. Reaction-diffusion model as a framework for understanding biological pattern formation. Science 329 (2010) 1616–1620. doi:10.1126/science.1179047.

4. Halatek J, Brauns F, Frey E. Self-organization principles of intracellular pattern formation. Philosophical Transactions of the Royal Society B: Biological Sciences 373 (2018) 20170107. doi:10.1098/rstb.2017.0107.

5. Schweisguth F, Corson F. Self-Organization in Pattern Formation. Developmental Cell 49 (2019) 659–677. doi:10.1016/j.devcel.2019.05.019.

6. Rietkerk M, van de Koppel J. Regular pattern formation in real ecosystems. Trends in Ecology & Evolution 23 (2008) 169–175. doi:10.1016/j.tree.2007.10.013.

7. Ramm B, Heermann T, Schwille P. The E. coli MinCDE system in the regulation of protein patterns and gradients. Cellular and Molecular Life Sciences 76 (2019) 4245–4273. doi:10.1007/s00018-019-03218-x.

8. Kretschmer S, Schwille P. Pattern formation on membranes and its role in bacterial cell division. Current Opinion in Cell Biology 38 (2016) 52–59. doi:10.1016/j.ceb.2016.02.005.

9. Mizuuchi K, Vecchiarelli AG. Mechanistic insights of the Min oscillator via cell-free reconstitution and imaging. Physical Biology 15 (2018) 031001. doi:10.1088/1478-3975/aa9e5e.

10. Wettmann L, Kruse K. The Min-protein oscillations in Escherichia coli : an example of self-organized cellular protein waves. Philosophical Transactions of the Royal Society B: Biological Sciences 373 (2018) 20170111. doi:10.1098/rstb.2017.0111.

11. Frey E, Halatek J, Kretschmer S, Schwille P. Protein Pattern Formation. Bassereau P, Sens P, editors, Physics of Biological Membranes (Cham: Springer International Publishing) (2018), 229–260. doi:10.1007/978-3-030-00630-310.

12. Vecchiarelli AG, Li M, Mizuuchi M, Mizuuchi K. Differential affinities of MinD and MinE to anionic phospholipid influence Min patterning dynamics in vitro: Flow and lipid composition effects on Min patterning. Molecular Microbiology 93 (2014) 453–463. doi:10.1111/mmi.12669.

13. Heermann T, Steiert F, Ramm B, Hundt N, Schwille P. Mass-sensitive particle tracking to elucidate the membrane-associated MinDE reaction cycle. Nature Methods 18 (2021) 1239–1246. doi:10.1038/s41592-021-01260-x.

14. Meinhardt H, de Boer PAJ. Pattern formation in Escherichia coli: A model for the pole-to-pole oscillations of Min proteins and the localization of the division site. Proceedings of the National Academy of Sciences 98 (2001) 14202–14207. doi:10.1073/pnas.251216598.

15. Kruse K, Howard M, Margolin W. An experimentalist’s guide to computational modelling of the Min system. Molecular Microbiology 63 (2007) 1279–1284. doi:10.1111/j.1365-2958.2007.05607.x.

16. Vecchiarelli AG, Li M, Mizuuchi M, Hwang LC, Seol Y, Neuman KC, et al. Membrane-bound MinDE complex acts as a toggle switch that drives Min oscillation coupled to cytoplasmic depletion of MinD. Proceedings of the National Academy of Sciences 113 (2016) E1479–E1488. doi:10.1073/pnas.1600644113.

17. Denk J, Kretschmer S, Halatek J, Hartl C, Schwille P, Frey E. MinE conformational switching confers robustness on self-organized Min protein patterns. Proceedings of the National Academy of Sciences 115 (2018) 4553–4558. doi:10.1073/pnas.1719801115.

18. Brauns F, Pawlik G, Halatek J, Kerssemakers J, Frey E, Dekker C. Bulk-surface coupling identifies the mechanistic connection between Min-protein patterns in vivo and in vitro. Nature Communications 12 (2021) 3312. doi:10.1038/s41467-021-23412-5.

19. Ivanov V, Mizuuchi K. Multiple modes of interconverting dynamic pattern formation by bacterial cell division proteins. Proceedings of the National Academy of Sciences 107 (2010) 8071–8078. doi:10.1073/pnas.0911036107.

20. Ramm B, Glock P, Mücksch J, Blumhardt P, García-Soriano DA, Heymann M, et al. The MinDE system is a generic spatial cue for membrane protein distribution in vitro. Nature Communications 9 (2018) 3942. doi:10.1038/s41467-018-06310-1.

21. Loose M, Fischer-Friedrich E, Herold C, Kruse K, Schwille P. Min protein patterns emerge from rapid rebinding and membrane interaction of MinE. Nature Structural & Molecular Biology 18 (2011) 577–583. doi:10.1038/nsmb.2037.

22. Zieske K, Schweizer J, Schwille P. Surface topology assisted alignment of Min protein waves. FEBS Letters 588 (2014) 2545–2549. doi:10.1016/j.febslet.2014.06.026.

23. Van Rossum G, Drake FL. Python 3 Reference Manual (Scotts Valley, CA: CreateSpace) (2009).

24. Vecchiarelli AG, Li M, Mizuuchi M, Ivanov V, Mizuuchi K. MinE recruits, stabilizes, releases, and inhibits MinD interactions with membrane to drive oscillation. preprint, Microbiology (2017). doi:10.1101/109637.

25. Ramm B, Goychuk A, Khmelinskaia A, Blumhardt P, Eto H, Ganzinger KA, et al. A diffusiophoretic mechanism for ATP-driven transport without motor proteins. Nature Physics 17 (2021) 850–858. doi:10.1038/s41567-021-01213-3.

26. Meindlhumer S, Brauns F, Finžgar J, Kerssemakers J, Dekker C, Frey E. Directing min protein patterns with advective bulk flow. BioRxiv (2021). doi:10.1101/2021.12.23.474007.

27. Caspi Y, Dekker C. Mapping out Min protein patterns in fully confined fluidic chambers. eLife 5 (2016) e19271. doi:10.7554/eLife.19271.

28. Würthner L, Brauns F, Pawlik G, Halatek J, Kerssemakers J, Dekker C, et al. Bridging scales in a multiscale pattern-forming system. 2111.12043 [nlin, physics:physics] (2021). ArXiv: 2111.12043.

29. Robertson C. Theory and practical recommendations for autocorrelation-based image correlation spectroscopy. Journal of Biomedical Optics 17 (2012) 080801. doi:10.1117/1.JBO.17.8.080801.

30. Horn BKP, Schunck BG. Determining optical flow. Artificial Intelligence 17 (1981) 185–203. doi:10.1016/0004-3702(81)90024-2.

31. Mesbah M. Gradient-based optical flow: a critical review. ISSPA ‘99. Proceedings of the Fifth International Symposium on Signal Processing and its Applications (IEEE Cat. No.99EX359) (Queensland Univ. Technol) (1999), vol. 1, 467–470. doi:10.1109/ISSPA.1999.818213.

32. [Dataset] Michael, Ilya. scivision/pyoptflow: update implementation, compare 3 implementations (2021). doi:10.5281/zenodo.5574844.

33. Vanag VK, Epstein IR. Pattern formation mechanisms in reaction-diffusion systems. The International Journal of Developmental Biology 53 (2009) 673–681. doi:10.1387/ijdb.072484vv.

