## Supplementary Information for "Quantitative analysis of surface wave patterns of Min proteins"

#### 1 METHODS

Plots were created using Python(1, 2) and assembled into figures using Inkscape(3). A few panels were prepared with Fiji(4). Example Min protein data was recorded using a spinning disc confocal microscope and processed as outlined in the main text. Unless specified otherwise, data for MinE is shown. Patterns are displayed in the colormap *viridis*, except for the dual-channel images shown in Fig. 4.

The Python(1) code created for this study as well as example data can be found in open repositories on GitHub (`https://github.com`) and Zenodo (`https://zenodo.org`).
Find the GitHub repository at `https://github.com/M-Sabrina/MinDE_analysis_2022`.
Find the second release at Zenodo at: `https://zenodo.org/record/6724666`. The content of this paper refers to this second release.
Find the latest release at Zenodo at `https://doi.org/10.5281/zenodo.6500098`.

In the readme-file, we provide instructions on how to set up a Python environment "min_analysis", including a package "min_analysis_tools". The folder "min_analysis_scripts" contains scripts that use the tools to perform analysis on single stacks or batch processing of all stacks within a folder.

We provide several Jupyter notebooks, all of which can be immediately launched and viewed in a Jupyter binder, such as:
`https://mybinder.org/v2/gh/M-Sabrina/MinDE_analysis_2022/HEAD`.

The repository contains two Jupyter notebooks starting with "DEMO", one for global and one for local analysis. These notebooks demonstrate in a step-by-step fashion how the analysis pipelines for autocorrelation analysis and local wave propagation detection work, using examples of simulated as well as real data. Snapshots of these Jupyter notebooks (for one out of several possible real examples) are provided as supplementary material in the appendix of this document. We recommend visiting the binder or installing the package for interactivity.

Further, the repository contains a notebook "Quickstart_Min_analysis.ipynb", which can directly be used to (remotely) perform analysis on a pre-cleaned single Min pattern data stack. Instructions are directly included in the notebooks.

Additionally, we provide a series of Supplementary notebooks, which can be used to repeat controls that we performed to test our analysis strategies:

- *Supplementary_crestpoint_bias.ipynb*: shows how accurately the originally determined crest points (from optical flow analysis) co-localize with local intensity maxima.
- *Supplementary_noise_sensitivity.ipynb*: shows effect of noise on accuracy of global methods and local velocity detection.
- *Supplementary_accuracy_global_spatial.ipynb*: shows effect of decreasing the image size on the accuracy of global autocorrelation analysis, with focus on the ratio of image size and spatial wavelength.
- *Supplementary_accuracy_global_temporal.ipynb*: shows effect of shortening the image stack along the temporal direction, with focus on the ratio of stack length and oscillation period.
- *Supplementary_accuracy_velocity.ipynb*: shows effect of decreasing the image size on the accuracy of local velocity analysis.
- *Supplementary_compare_temporal_autocorrelation_methods.ipynb*: shows a comparison of reslicing-based temporal autocorrelation analysis (as presented in the paper) and pixelwise autocorrelation. An analysis performed on the dataset represented in Fig. 1 run on the Jupyter binder [24-06-2022] returned comparable results for the oscillation period within 0.2 s for the reslicing method and 17.4 s for the pixelwise method.

#### REFERENCES


**1** .Van Rossum G, Drake FL. *Python 3 Reference Manual* (Scotts Valley, CA: CreateSpace) (2009).

**2** .Hunter JD. Matplotlib: A 2d graphics environment. *Computing in science & engineering* **9** (2007) 90–95.

**3** .Inkscape Project. Inkscape. *https://inkscape.org* (2021).

**4** .Schindelin J, Arganda-Carreras I, Frise E, Kaynig V, Longair M, Pietzsch T, et al. Fiji: an open-source platform for biological-image analysis. *Nature Methods* **9** (2012) 676–682. doi:10.1038/nmeth.2019.

#### 2 SUPPLEMENTARY FIGURES

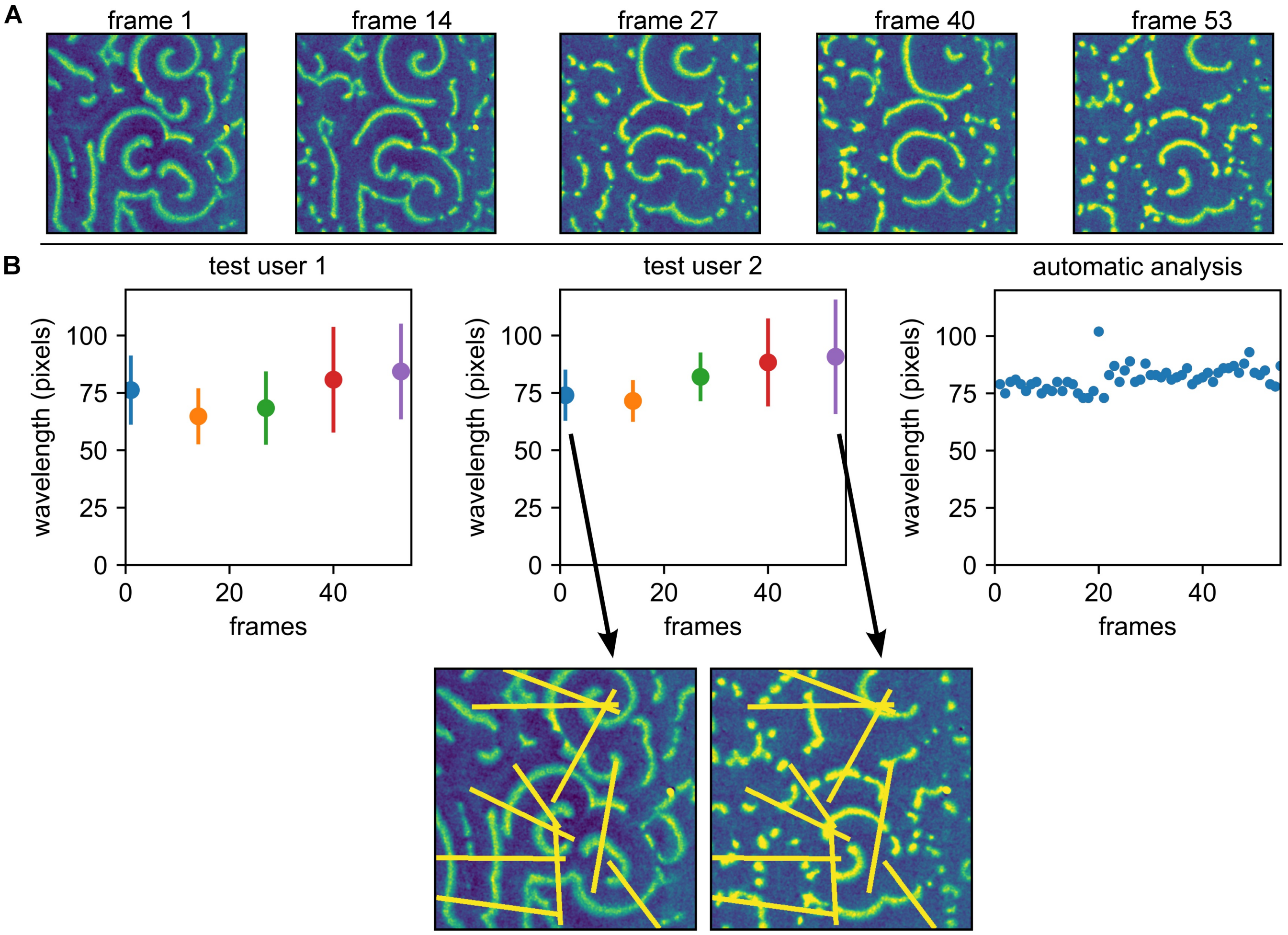


**Figure S1.** **A** Example frames from an image stack comprising 55 sequential frames of size 512 x 512 pixels). **B** For comparing the outcome of our automatized spatial autocorrelation routine to manual quantification, we asked two test persons to analyse the stack using Fiji(4) via the following procedure: Based upon the pattern presented in frame 1, pick ten cross-sections of line-width 10 pixels, which will be kept constant over the movie. For each cross-section, identify two peaks (if possible), then plot a line profile and identify the spacing of two consecutive peaks. This spacing is indicative of the wavelength. Perform this for frames 1, 14, 27, 40, and 53. Left and center panel: Manual results from test persons 1 and 2. Error bars denote standard deviations. Bottom panel shows the cross-sections selected by test user 2 for frames 1 and 53. Right panel: Results from the automated spatial correlation analysis.

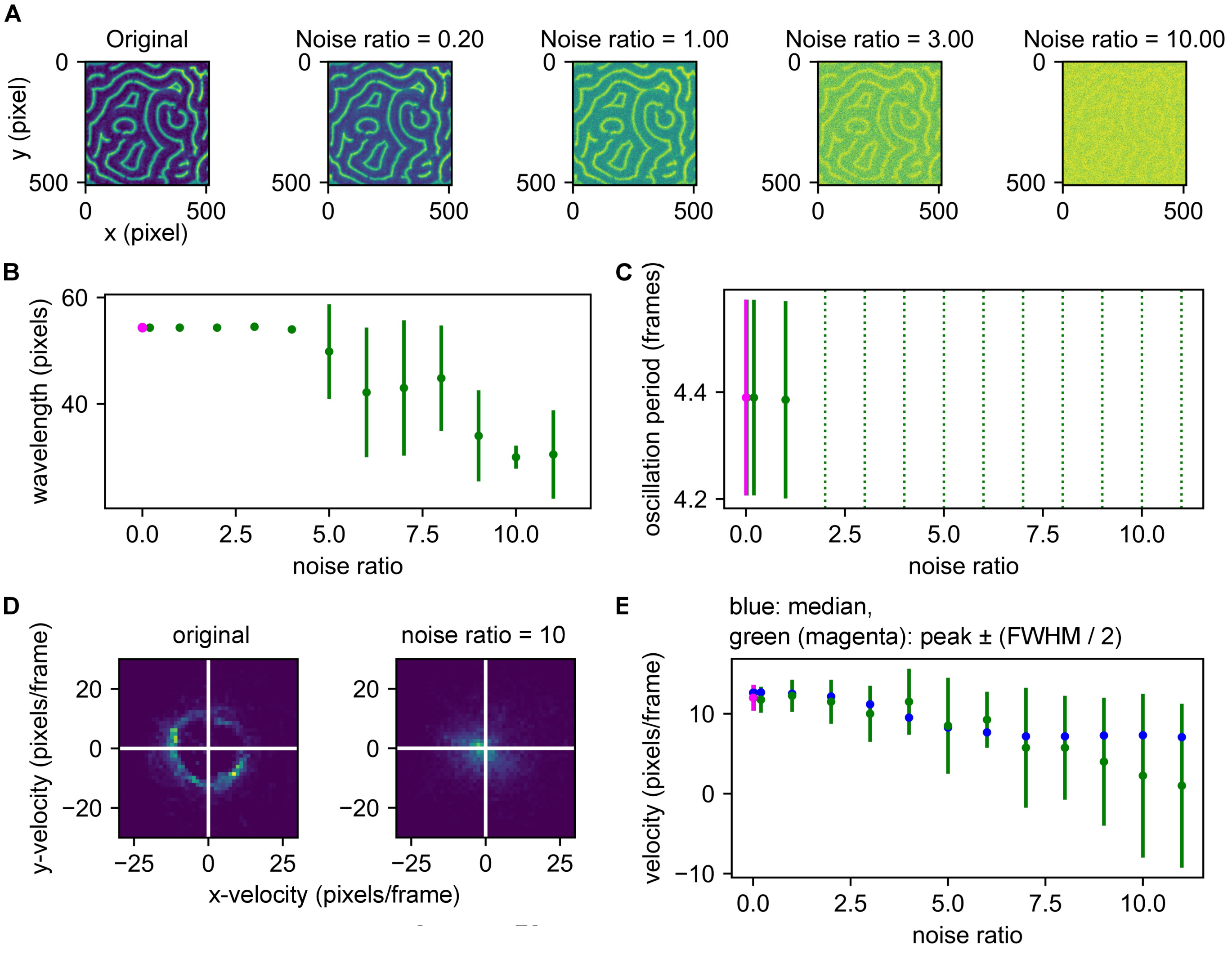


**Figure S2.** **A** Starting from a clean image stack comprising 20 sequential frames, we created a set of image stacks of decreasing quality by adding higher levels of Gaussian noise. Example frames of different noise ratios are shown here. Noise ratio refers to the standard deviation of added Gaussian noise with respect to the intrinsic image standard deviation. **B** Effect of noise on global spatial analysis, from analysis of first 6 frames. Resulting wavelength and standard deviation without noise in magenta, for increasing noise ratios in green. **C** Effect of noise on global temporal analysis from analysis on 20 frames, for 10 horizontal and vertical line traces each. Resulting period and standard deviation without noise in magenta, for increasing noise ratios in green. Green dashed lines indicated noise ratios for which our algorithm failed to return a result. **D** Example 2D histograms showing the effect of strong noise on local velocity analysis. **E** Results for local velocity analysis on first 6 frames. Resulting peak velocity magnitude and standard deviation (from FWHM of distribution) without noise in magenta, for increasing noise ratios in green. Median velocities in blue. *Also see notebook Supplementary_noise_sensitivity.ipynb*

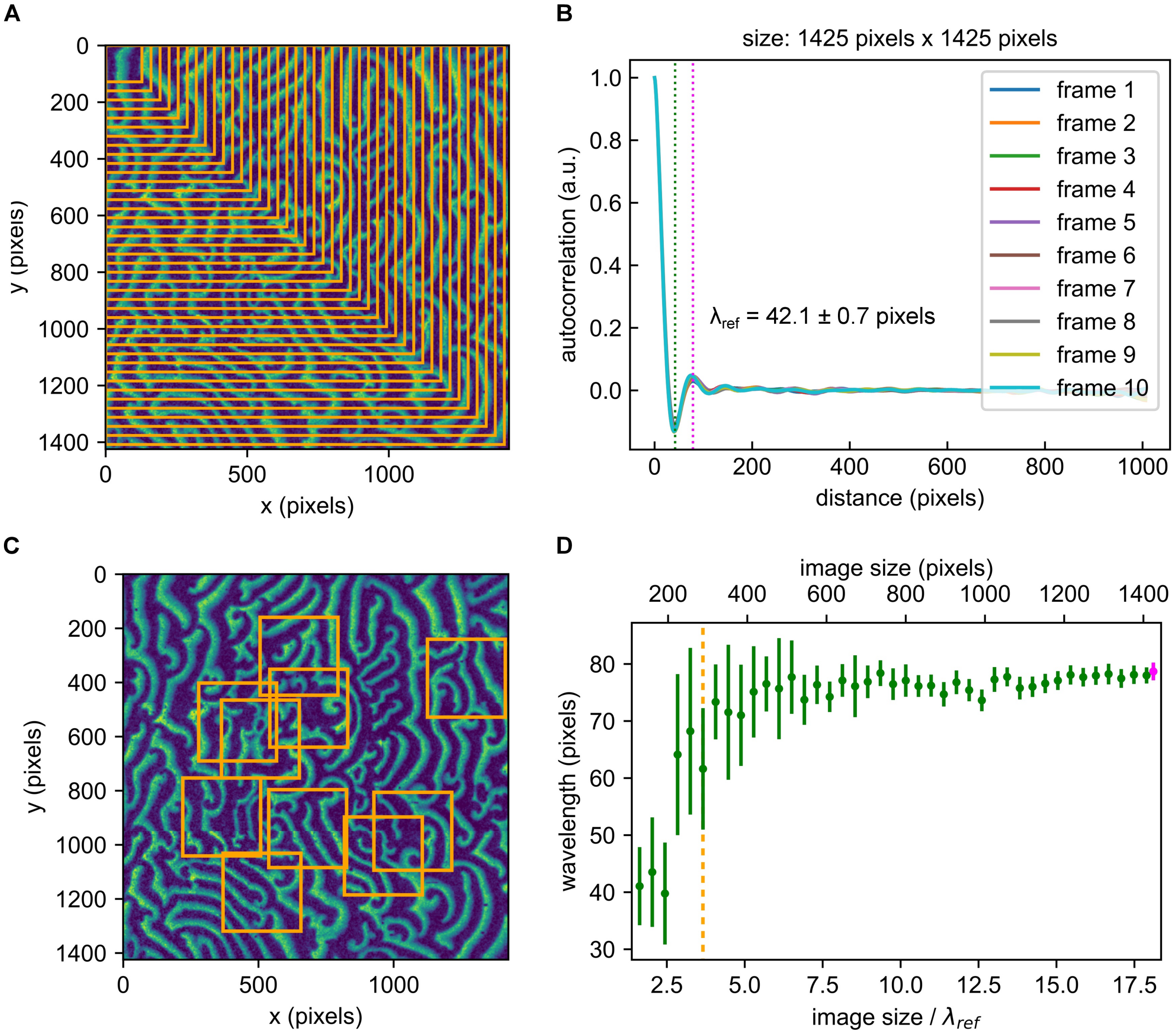


**Figure S3.** **A** Image stack comprising 10 consecutive frames, sequentially cropped to smaller sizes. **B** Spatial autocorrelation results for full-scale image stack. **C** For each cropping sizes, 10 random surface locations are chosen. **D** Spatial autocorrelation results for different cropping sizes, averaged from 10 randomly selected surface regions such as shown in C. Resulting wavelength and standard deviation for full in magenta (referred to as $\lambda_{ref}$), for cropped images in green. $x$-axis denotes both absolute image size (in pixels) as well as image size with respect to $\lambda_{ref}$. Image size for the example in C indicated by orange dashed line. *Also see notebook Supplementary_accuracy_global_spatial.ipynb*

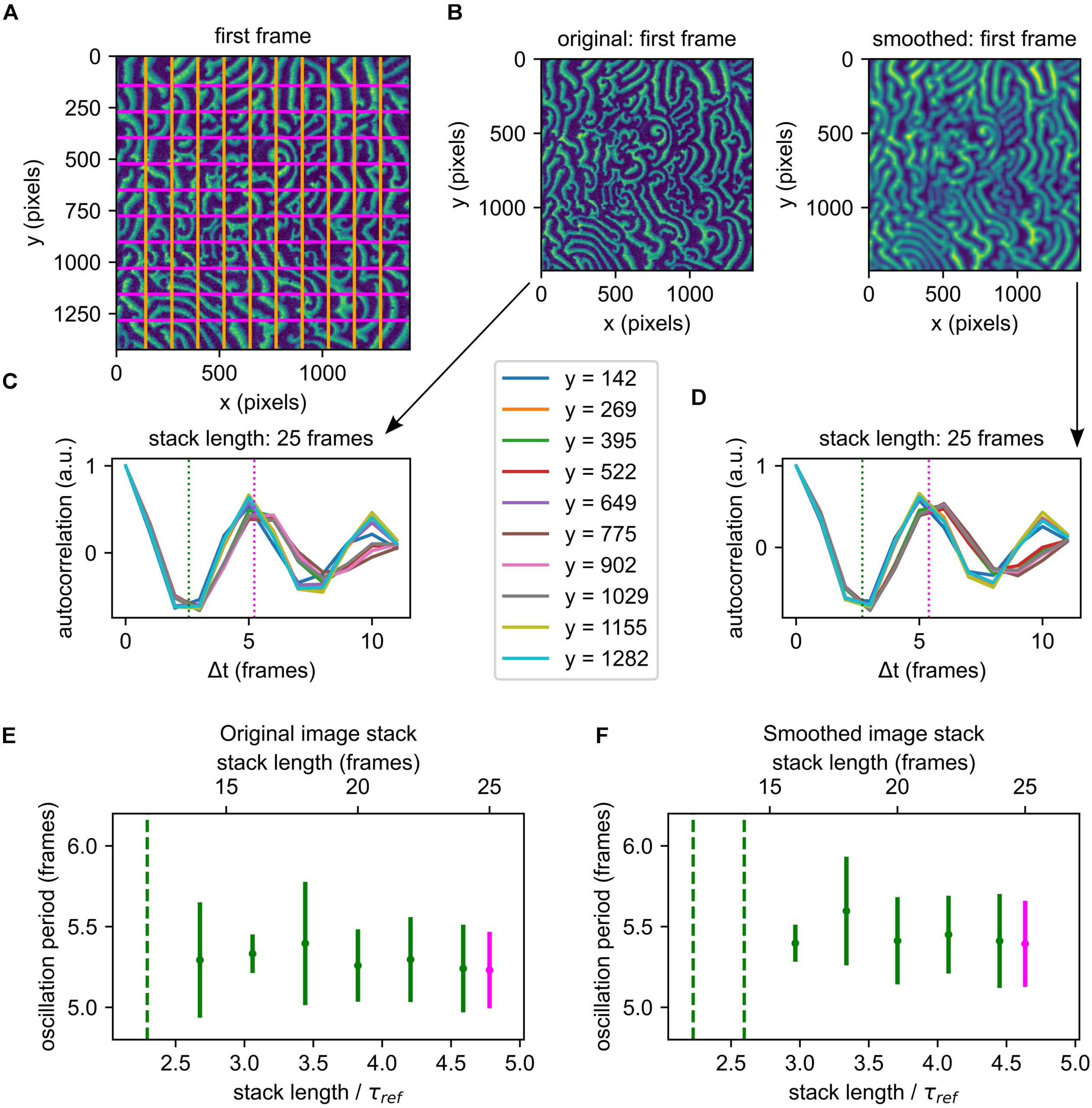


**Figure S4.** Starting from an image stack comprising 25 frames, we sequentially crop it in the temporal direction and study the effect of this shortening on both an unsmoothed and smoothed image. **A** Analysis is performed along the horizontal and vertical lines indicated. Each line (corresponding to one resliced image stack) provides a result from global temporal autocorrelation analysis. These results are collected to obtain a mean and standard deviation for the oscillation period. **B** Original (unsmoothed) and smoothed image stacks, first frame shown for each. **C** Full analysis (25 frames) for original image stack as shown in B, results providing the reference oscillation period $\tau_{ref}$. **D** Full analysis (25 frames) on strongly smoothed image stack as shown in B, results providing the reference oscillation period $\tau_{ref}$. **E** Results for original image stack shown as dependent on absolute stack length (in frames) and with respect to $\tau_{ref}$. Resulting period and standard deviation for full stack in magenta (referred to as $\tau_{ref}$), for cropped image stacks in green. Green dashed lines indicated stack lengths at which our algorithm failed to return a result. **F** Same as E for strongly smoothed stack. *Also see notebook Supplementary_accuracy_global_temporal.ipynb*

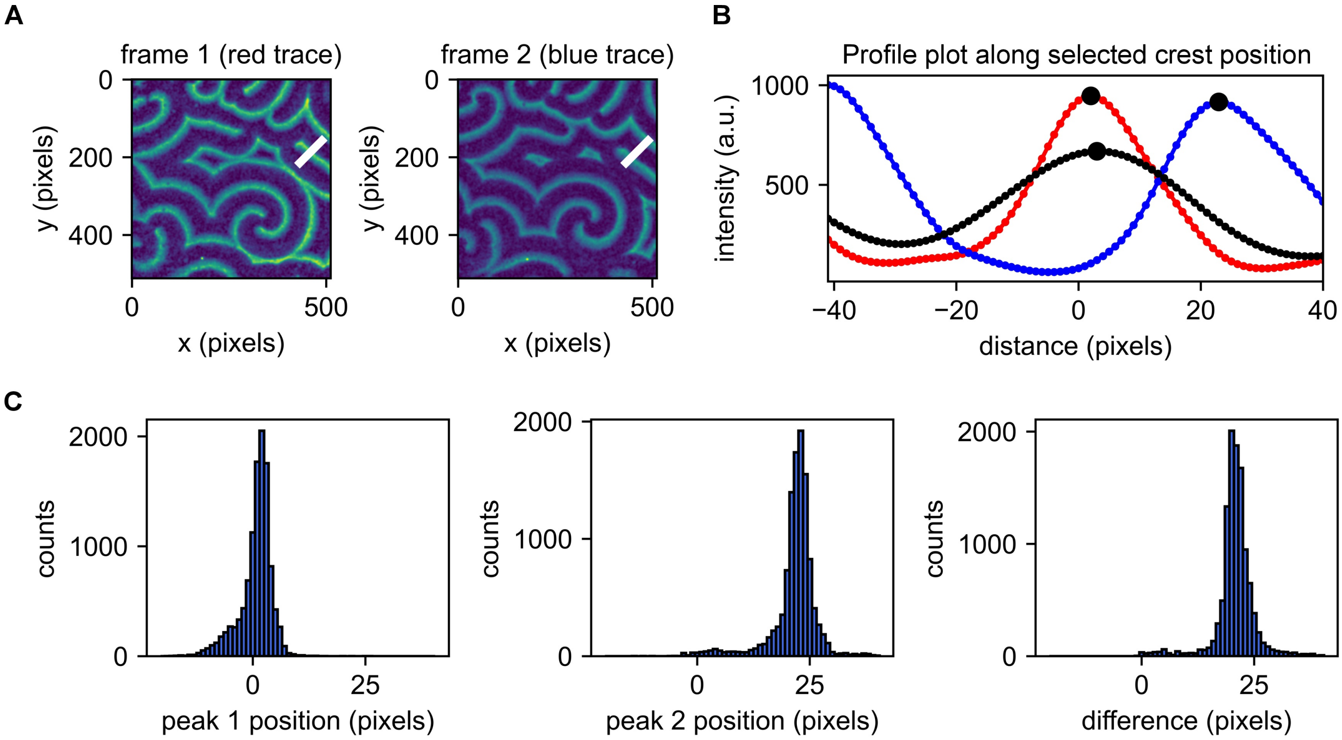


| kernel | peak position 1 | peak 1 FWHM | peak shift | peak shift FWHM |
|---|---|---|---|---|
| 10 | 1.5 | 2 | 19.5 | 6 |
| 20 | 1.5 | 3 | 19.5 | 5 |
| 30 | 1.5 | 3 | 19.5 | 5 |
| 40 | 1.5 | 4 | 19.5 | 6 |
| 50 | 2.5 | 7 | 19.5 | 6 |
| 60 | 1.5 | 19 | 20.5 | 5 |

**Figure S5.** **A** Example frames of a sequential image series. **B** Example intensity traces of frames 1 (red), frame 2 (blue) and smoothed intensity trace of frame 1 (black, smoothing kernel 35 pixels) at the location indicated in A. **C** Histograms showing locations of peaks from frame 1 (peak 1 position) and frame 2 (peak 2 position) with respect to the originally assumed crest position as determined from optical flow analysis.**Table:** Position of peak 1 (pixels) from originally determined crest position as dependent on smoothing kernel used for optical flow analysis. *Also see notebook Supplementary_crestpoint_bias.ipynb*

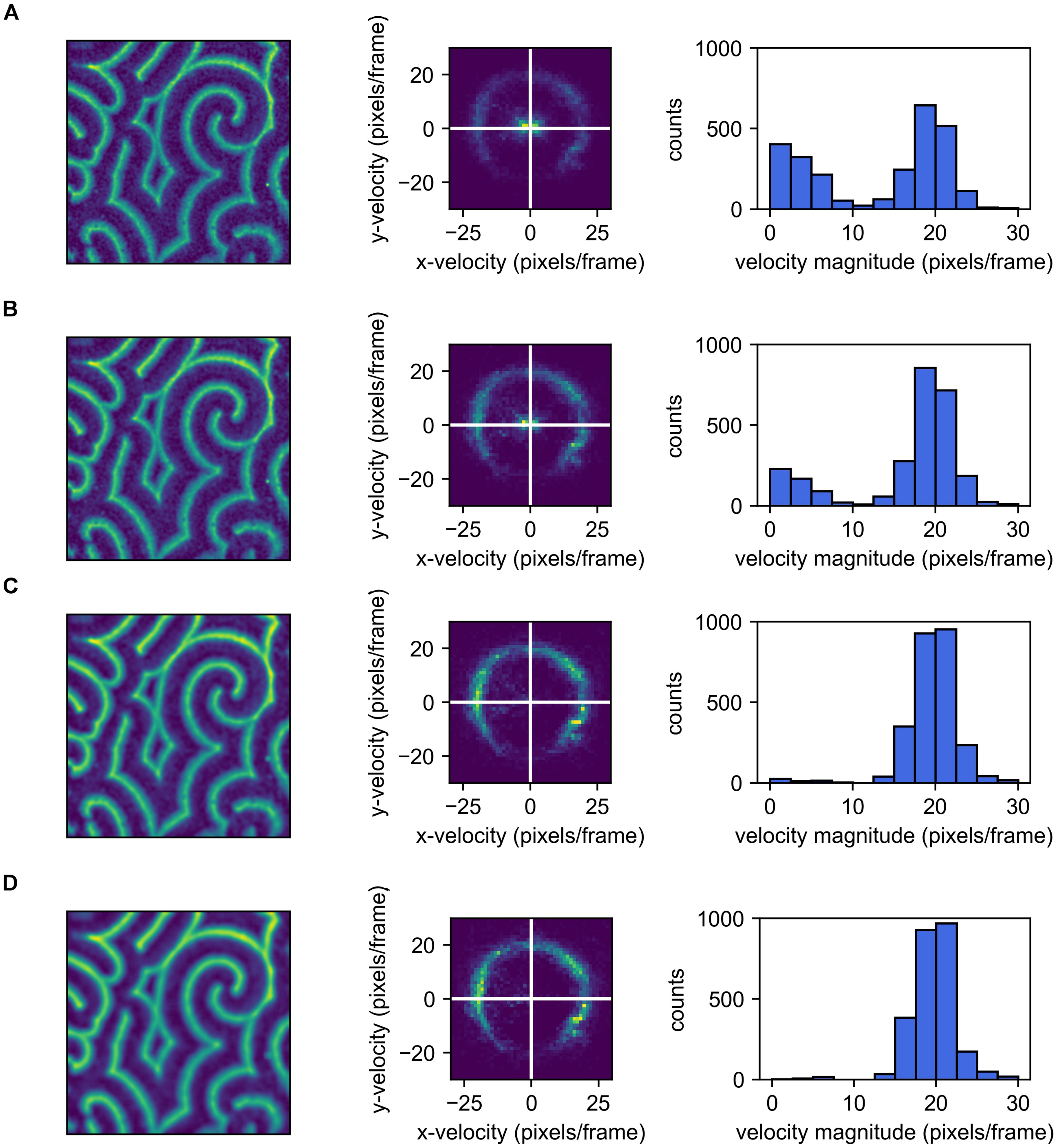


**Figure S6.** Effect of initial image smoothing on the quantification of local velocity (with for the optical flow analysis, a constant additional smoothing step with kernel of 35 pixels size). Quantification was performed over 10 consecutive frames of an example image stack, image size (512 x 512 pixels). **A** No smoothing. **B** Smoothing kernel of 5 pixels size. **C** Smoothing kernel of 10 pixels size. **D** Smoothing kernel of 15 pixels size.

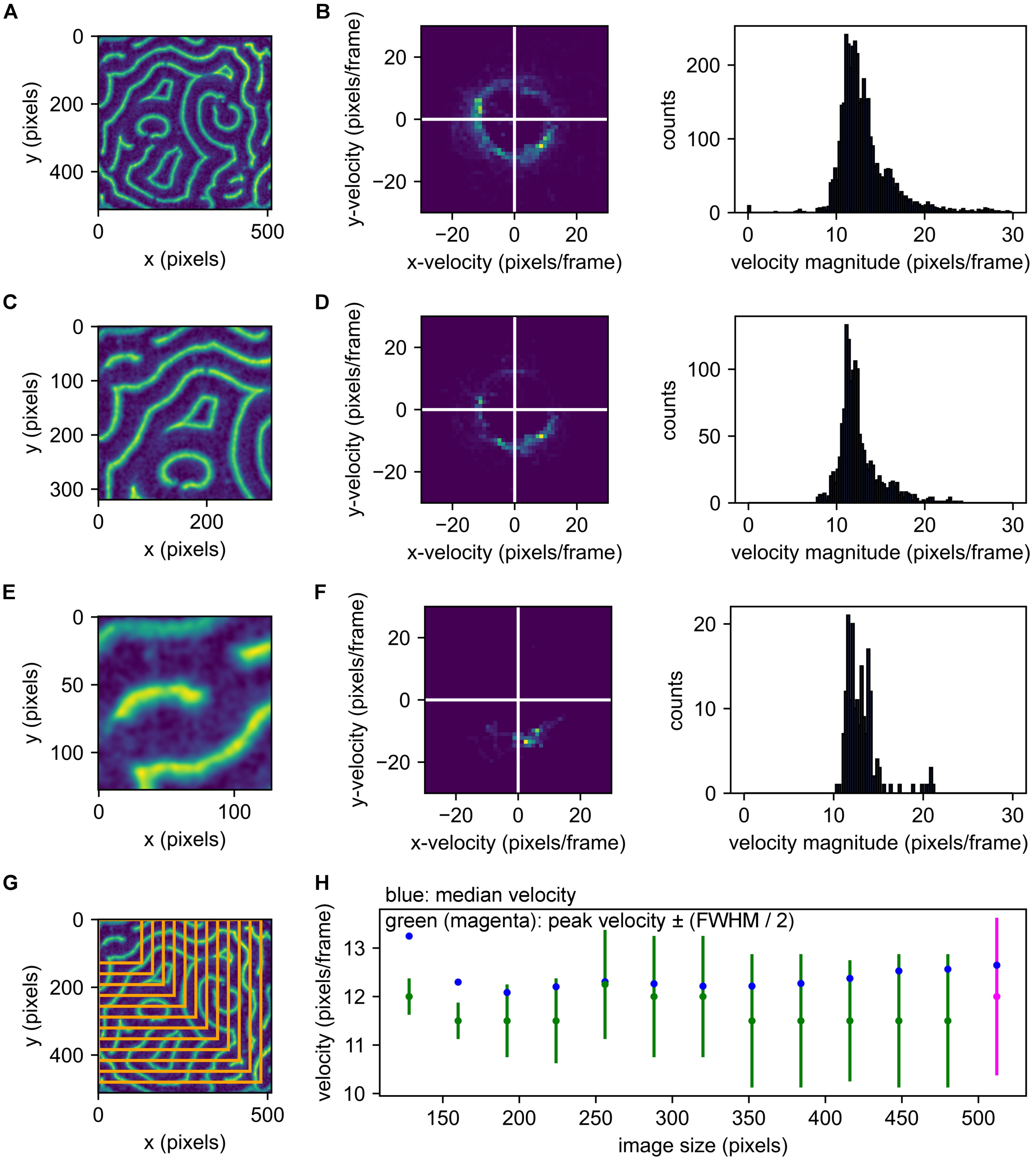


**Figure S7.** Effect of image size on local velocity analysis, as shown on an example image stack comprising 4 consecutive frames, sequentially cropped to smaller sizes. **A** Original image size, first frame. **B** Local velocity analysis results for original image size. **C** and **D** Same as A and B for cropped image size. **E** and **F** Same as A and B for further cropped image. **G** Illustration of all used cropping sizes. **H** Results for local velocity analysis on 4 frames for cropping sizes illustrated in G. Resulting peak velocity magnitude and standard deviation (from FWHM of distribution) for full image size in magenta, for cropped images in green. Median velocities in blue. *Also see notebook Supplementary_accuracy_velocity.ipynb*

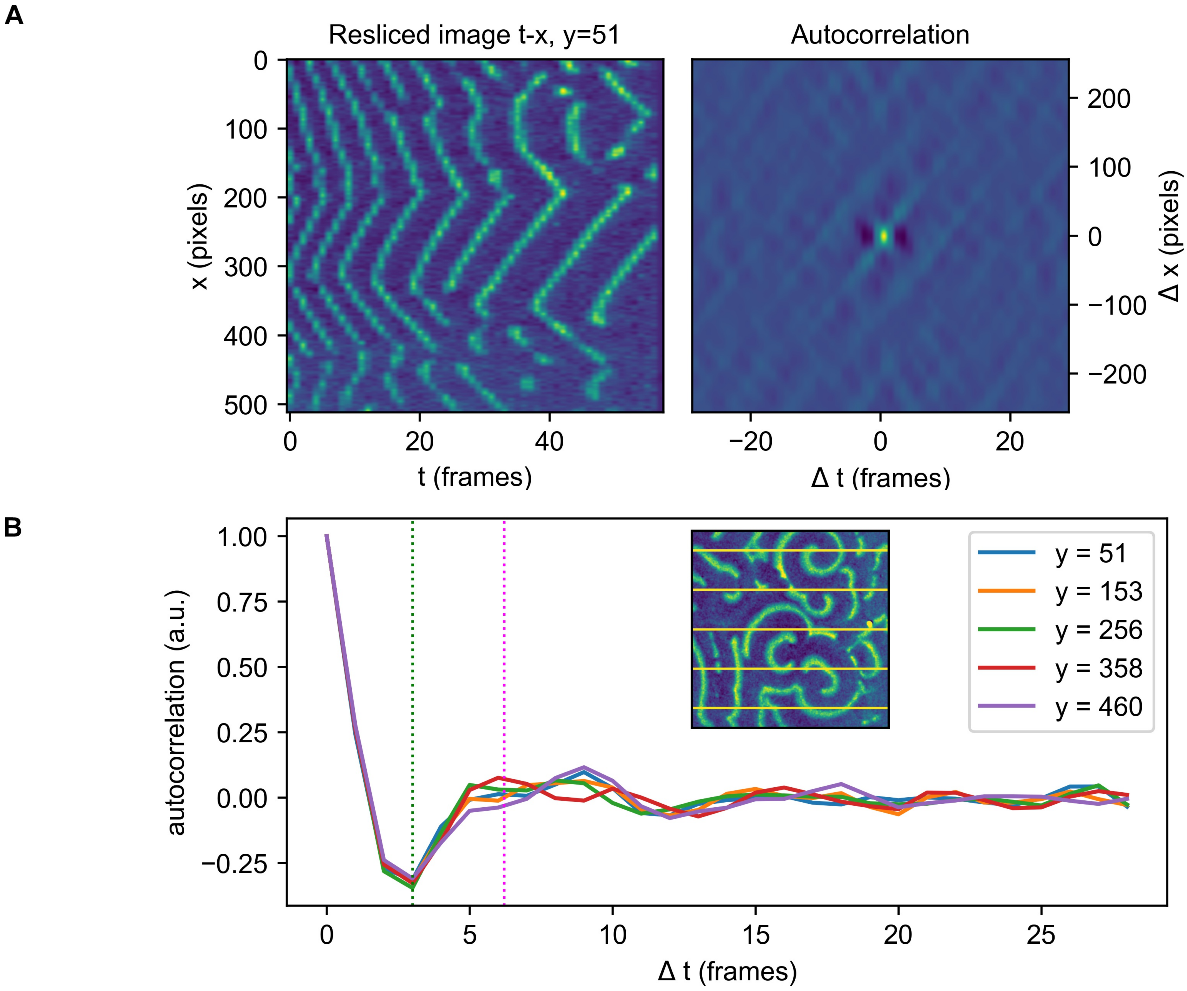


**Figure S8.** Temporal autocorrelation analysis performed for the image stack shown in Fig. S1A. **A** Resliced image (set of kymographs) for fixed spatial $y$-coordinate (left) and autocorrelation map (right). **B** $\Delta x = 0$ traces for fixed $y$-coordinates. Inset shows fixed $y$-positions along which analysis was performed superimposed on frame 1. The magenta dashed line indicates the average position of the first maximum, which in this case yields an incorrect result for the oscillation period.

---

### APPENDIX

### DEMO_MinDE_global_analysis

June 24, 2022

#### 1 Wavelength and oscillation period of MinDE patterns

*Jacob Kerssemakers, Sabrina Meindlhumer, Cees Dekker lab, 2022*

The flowing, semi-periodic patterns of MinDE pose some challenges to quantification. Here, we illustrate a systematic approach to extract global parameters, in particular the global wavelength $\mu$ and global oscillation period $\tau$.

**READ-ME:** Cells in this notebook need to be executed sequentially. Upon starting to explore this notebook, click the double-arrow symbol above (*Restart the kernel, then re-run the whole notebook*) and hit "Restart" to ensure all required packages are loaded. After that, the notebook will take a few moments to set up, and figures/plots will re-appear one by one. At distinct positions in the notebook, the user is invited to change numeric input. After doing so, the notebook needs to be executed anew at least from this point on for changes to be applied. The notebook can be re-run from any given point onwards by clicking *Run* in the menu-bar above, and selecting *Run Selected Cell and All Below*. Alternatively, the double-arrow symbol can be used again, which will re-run the notebook from the start. This will take a moment longer, but will have the same effect.

##### 1.1 Setup

Import of standard modules and assisting custom-made modules. Needs to be executed at least once to ensure correction functionality (see instructions above).

[1]:
```
import numpy as np
from skimage import io
import matplotlib.pyplot as plt
from pathlib import Path

from min_analysis_tools import correlation_tools
from min_analysis_tools import get_data

# Reload modules automatically before executing code
%reload_ext autoreload
%autoreload 2

# Figure quality
import matplotlib as mpl
mpl.rcParams['figure.dpi'] = 300
```

#### 1.2 Select example

Choose an example from the provided stacks in the list below: (1) Simulated spiral (2) Min spiral (example data) (3) Min southeast-directed traveling waves (example data) (4) Min west-directed traveling waves (example data) (5) Min large stitched pattern (example data) (6) Min horizontally stitched pattern (example data) Choose the example by setting the variabel *selection* in the code-box below. The notebook needs to be re-run (at least from this point onwards) for changes to be applied.

[2]:
```
selection = 2  # set to 1 ... 6
```

[3]:
```
if selection not in np.arange(1, 7):
    print("Invalid selection. Set to selection 1 (Simulated spiral).")
    selection = 1
    MinDE_st = get_data.generate_pattern(
        lambda_t=20, lambda_x=1, size=512, N_frames=50, demo=False
    )
elif selection == 1:  # "Simulated spiral"
    MinDE_st = get_data.generate_pattern(
        lambda_t=20, lambda_x=1, size=512, N_frames=50, demo=False
    )
elif selection > 1:
    if selection == 2:  # "Min spiral (example data)"
        stack_path = Path().cwd() / "example_data" / "demo_spiral.tif"
    if selection == 3:  # "Min southeast-directed traveling waves (example data)"
        stack_path = Path().cwd() / "example_data" / "demo_southeast.tif"
    if selection == 4:  # "Min west-directed traveling waves (example data)"
        stack_path = Path().cwd() / "example_data" / "paper_west_E.tif"
    if selection == 5:  # "Min large stitched pattern (example data)"
        stack_path = Path().cwd() / "example_data" / "demo_square_stitch.tif"
    if selection == 6:  # "Min horizontally stitched pattern (example data)"
        stack_path = Path().cwd() / "example_data" / "demo_horizontal_stitch.tif"
    MinDE_st = io.imread(stack_path)
nt, ny, nx = np.shape(MinDE_st)

fig, ax = plt.subplots()
ax.imshow(MinDE_st[0, :, :])
ax.set_xlabel("x (pixels)")
ax.set_ylabel("y (pixels)")
plt.show()

print(f"Current selection: {selection}")
```

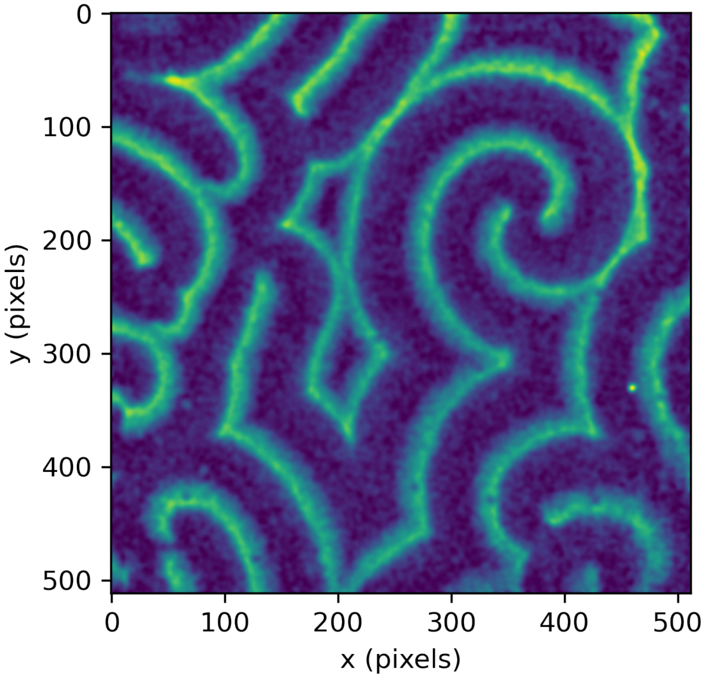


```
Current selection: 2
```

##### 1.3 Spatial autocorrelation

Spatial autocorrelation analysis is performed on a set number of images per movie. For each autocorrelation output image, a radial profile is recorded starting from the main central correlation peak. The resulting spatial radial correlation curves are subjected to maxima analysis. The first maximum after radius $R = 0$ indicates the most predominant distance between wave edges, irrespective of propagation direction. This distance is identified as the pattern's global wavelength. First, we perform autocorrelation on a set number of frames of our loaded stack. For each analyzed frame, we obtain a correlation matrix such as the one below. You can change this number in the code-box below. The notebook needs to be re-run (at least from this point on) for changes to apply.

[4]:
```
frames_to_analyse = 10  # integer number, default: 10
```

[5]:
```
# calculate spatial autocorrelation maps for all analysed frames
(
    crmx_storage,
    fig,
    ax_corr,
    ax_orig,
) = correlation_tools.get_spatial_correlation_matrixes(
    MinDE_st, frames_to_analyse, demo=True
)
```

```
Analysing 10 frames
```

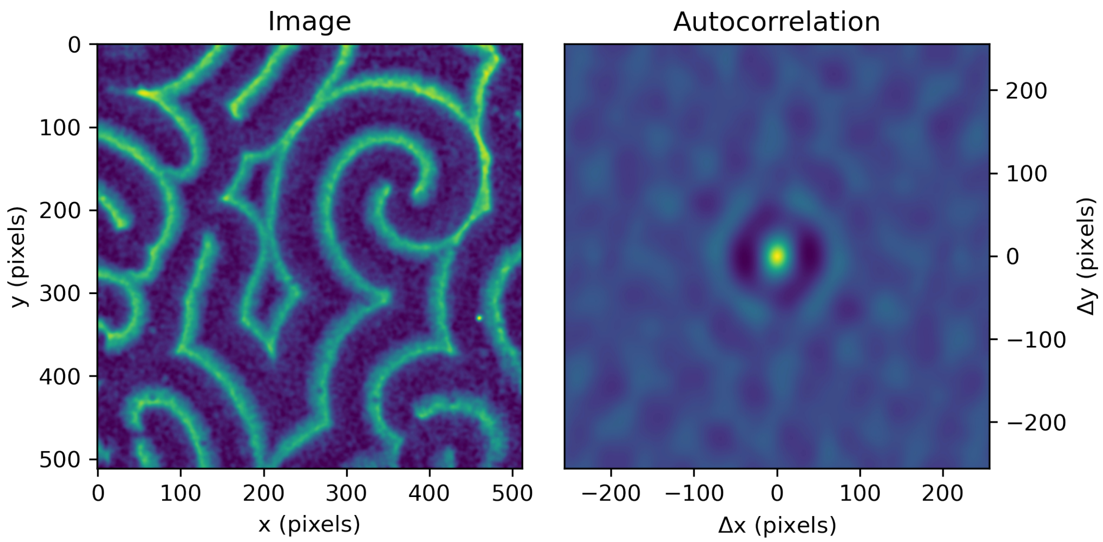


#### 1.4 Radial profile traces

Next, we perform radial averaging starting from the center peak. Ideally, we obtain a curve showing a clear first peak, indicating high correlation. We interpret this peak as representing the pattern's predominant, "global" wavelength $\mu$.

[6]:
```
# calculate radially averaged profile traces and analyze them with respect to␣
↪first min and max
(
    min_pos,
    min_val,
    max_pos,
    max_val,
    fig,
    ax,
) = correlation_tools.analyze_radial_profiles(crmx_storage, demo=True)
```

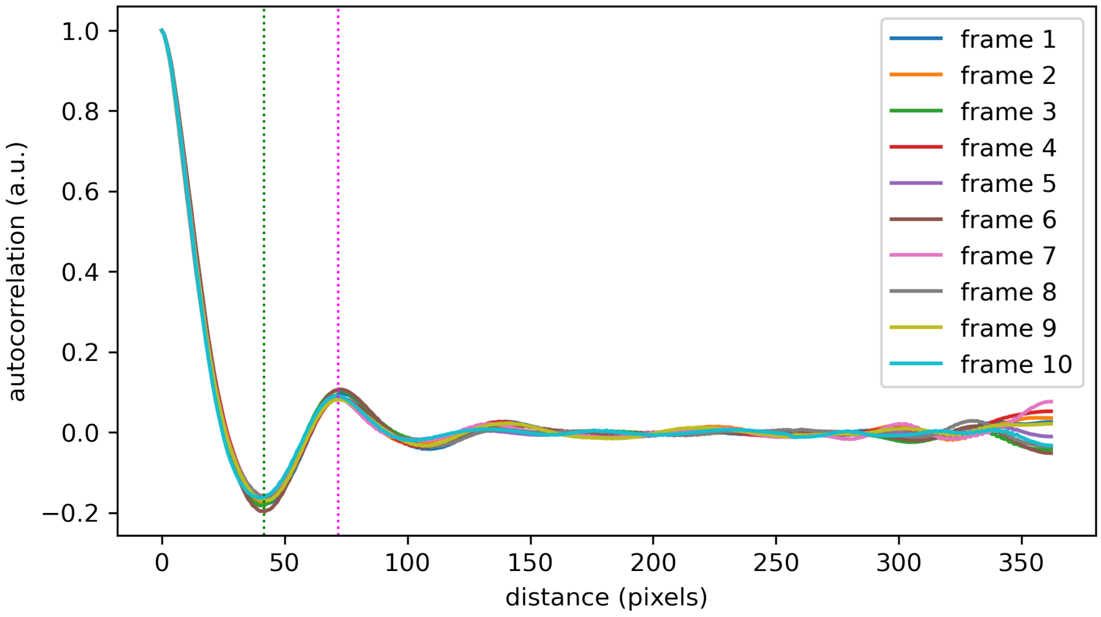


Wavelengths obtained from all analysed frames can be averaged:

[7]:
```
# output characteristic parameters
print(f"Mean position of first valley: {np.mean(min_pos):.02f}")
print(f"Mean position of first peak: {np.mean(max_pos):.02f} (wavelength in␣
 ↪pixels)")
```

```
Mean position of first valley: 41.60
Mean position of first peak: 71.90 (wavelength in pixels)
```

##### 1.5 Temporal autocorrelation

For temporal correlation, we generate a set of $t$-$x$ or $t$-$y$ kymographs per movie, evenly distributed over a set middle fraction fraction of an image. For each set of kymographs, autocorrelation analysis is performed. The $\Delta x$=0 or $\Delta y$=0 line of the resulting autocorrelation maps then in effect represent a temporal correlation curve averaged over all the original image points on this line. In other words, they represent average temporal correlation signals sampled from all selected surface locations. Analogous to the spatial correlation analysis, the first maximum after $\Delta t = 0$ indicates a main oscillation period. We start by reslicing our images in $x$- or $y$-direction. Note that one frame in such a stack then corresponds to one horizontal or vertical cross-section in the original imaged region, meaning that the frames in our stack are comprised of kymographs of all points along this line.

[8]:
```
# reslice frames
MinDE_shift_tx = np.moveaxis(MinDE_st, 0, -1)  # creates t-x resliced frames
MinDE_shift_yt = np.moveaxis(MinDE_st, -1, 0)  # creates y-t resliced frames
```

```
# next transpose y-t slices to have t axis in 1st dimension
MinDE_shift_ty = np.empty((nx, ny, nt))
for frame in range(nx):
    MinDE_shift_ty[frame, :, :] = np.transpose(MinDE_shift_yt[frame, :, :])
```

Using these resliced stacks makes it easier to access the temporal information within our data. We will now perform autocorrelation for selected frames of our resliced stacks, corresponding to cross sections along the $x$- or $y$-axis along our original images as shown above. The $x$- and $y$-positions of these lines are determined by the two parameters *reps_per_kymostack* and *kymoband*. These input numbers can be changed in the code-box below. The notebook needs to be re-run (at least from this point on) for changes to apply.

[9]:
```
reps_per_kymostack = 5  # pick ... kymographs around middle (integer number,␣
 ↪default: 5)
kymoband = 0.8  # analyse middle ... part of image (< 1, default: 0.8)
```

Below, the colored lines indicate all cross-sections for constant $y$ along which autocorrelation is performed, each corresponding to one particular frame in the $t$-$x$ resliced stack. The thick, red lines highlights the first horizontal cross-section at constant $y$, for which the exemplary $t$-$x$ slice and its autocorrelation map are shown. Note that the slice is comprised of a series of kymographs for all $x$ positions along that particular constant $y$.

[10]:
```
print(f"Current reps_per_kymostack: {reps_per_kymostack}")
print(f"Current kymoband: {kymoband}")

slices2analyze_x = ny * np.linspace(
    0.5 - kymoband / 2, 0.5 + kymoband / 2, reps_per_kymostack
)
slices2analyze_x = slices2analyze_x.astype(int)

fig, ax = plt.subplots(1, 1)
ax.imshow(MinDE_st[0, :, :])
ax.set_xlabel("x (pixels)")
ax.set_ylabel("y (pixels)")
ax.set_title("original image (first frame)")
for num_y, y_axis in enumerate(slices2analyze_x):
    if num_y == 0:
        ax.axhline(y=y_axis, color="red", linewidth=5)
    else:
        ax.axhline(y=y_axis, color="magenta")
```

```
Current reps_per_kymostack: 5
Current kymoband: 0.8
```

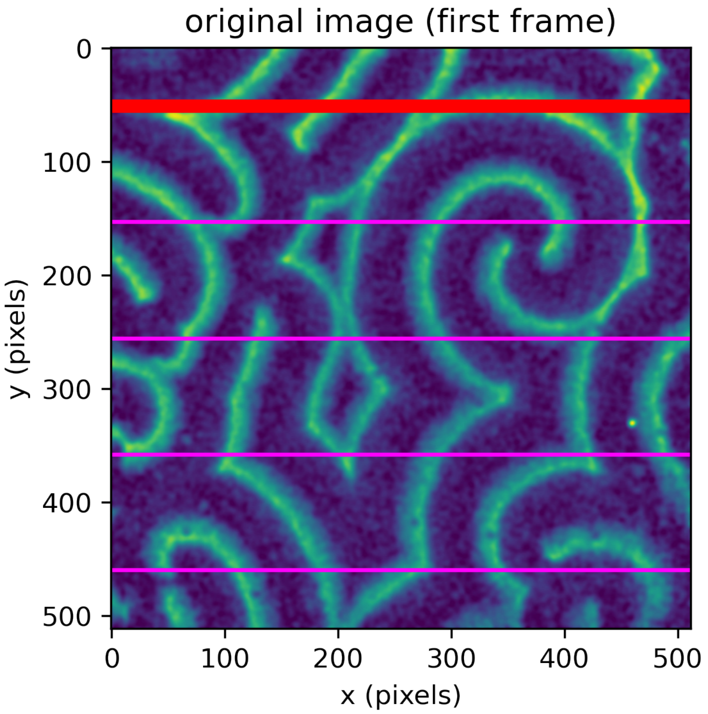


[11]:
```
selection_reslice_x = f"{selection}_x_resliced"
(
    crmx_storage_x,
    slices2analyze,
    fig,
    ax_corr,
    ax_orig,
) = correlation_tools.get_temporal_correlation_matrixes(
    MinDE_shift_tx,
    "x",
    kymoband,
    reps_per_kymostack,
    demo=True,
)
```

```
Analyzing t-x slices for y = [ 51 153 256 358 460]
```

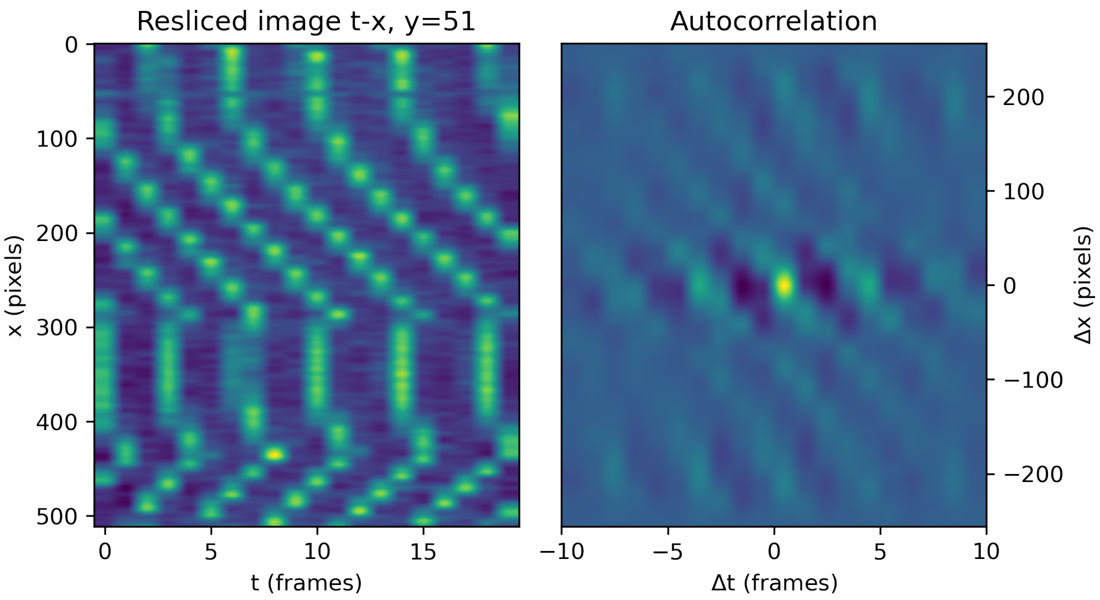


The $\Delta x = 0$ line of these autocorrelation maps then in effect represents a temporal correlation curve averaged over all the original image points along this cross-section at constant $y$. Analogous to the spatial correlation analysis, the first maximum after $\Delta t = 0$ indicates a main oscillation period. As temporal autocorrelation curves will generally be more coarse as they are not obtained from radial averaging, we perform cubic spline fitting and determine the peak and valley from extreme point analysis.

[12]:

```
# analyse selected traces with respect to first min and max
(
    first_min_pos_x,
    first_min_val_x,
    first_max_pos_x,
    first_max_val_x,
    fig,
    ax,
) = correlation_tools.analyze_temporal_profiles(
    "x", crmx_storage_x, slices2analyze_x, demo=True
)
```

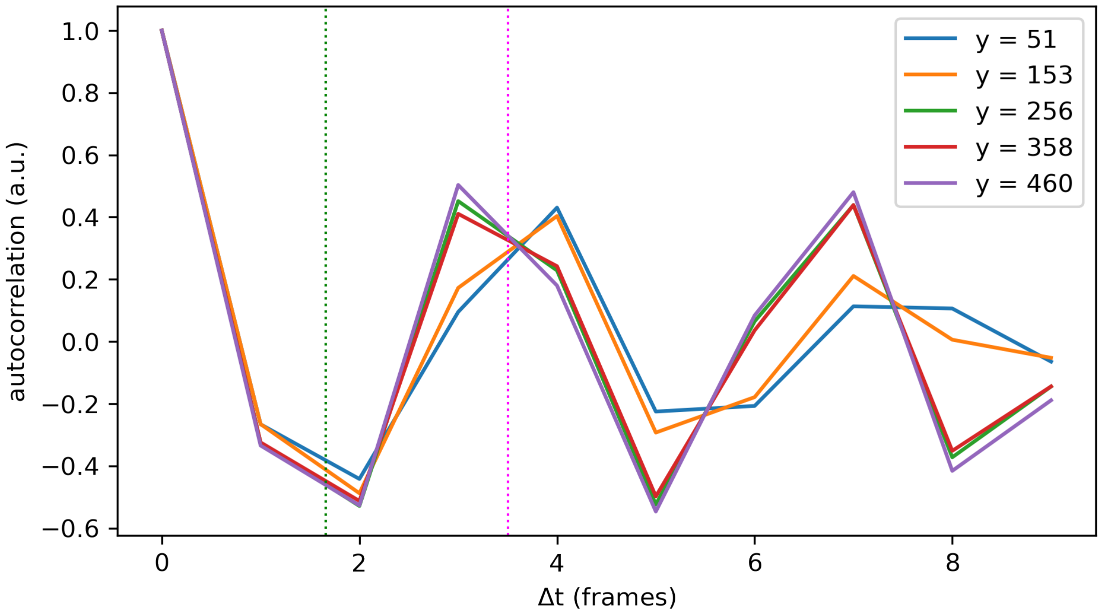


The same analysis pipeline can be performed for cross-sections at constant $x$. Below, the colored lines indicate all cross-sections for constant $x$ along which autocorrelation is performed, each corresponding to one particular frame in the $t$-$y$ resliced stack. The thick, red lines highlights the first horizontal cross-section at constant $x$, for which the exemplary $t$-$y$ slice and its autocorrelation map are shown. Again, note that the slice is comprised of a series of kymographs for all $y$ positions along that particular constant $x$.

[13]:
```
print(f"Current reps_per_kymostack: {reps_per_kymostack}")
print(f"Current kymoband: {kymoband}")

slices2analyze_y = nx * np.linspace(
    0.5 - kymoband / 2, 0.5 + kymoband / 2, reps_per_kymostack
)
slices2analyze_y = slices2analyze_y.astype(int)

fig, ax = plt.subplots(1, 1)
ax.imshow(MinDE_st[0, :, :])
ax.set_xlabel("x (pixels)")
ax.set_ylabel("y (pixels)")
ax.set_title("original image (first frame)")
for num_x, x_axis in enumerate(slices2analyze_y):
    if num_x == 0:
        ax.axvline(x=x_axis, color="red", linewidth=5)
    else:
        ax.axvline(x=x_axis, color="magenta")
```

```
Current reps_per_kymostack: 5
```

```
Current kymoband: 0.8
```

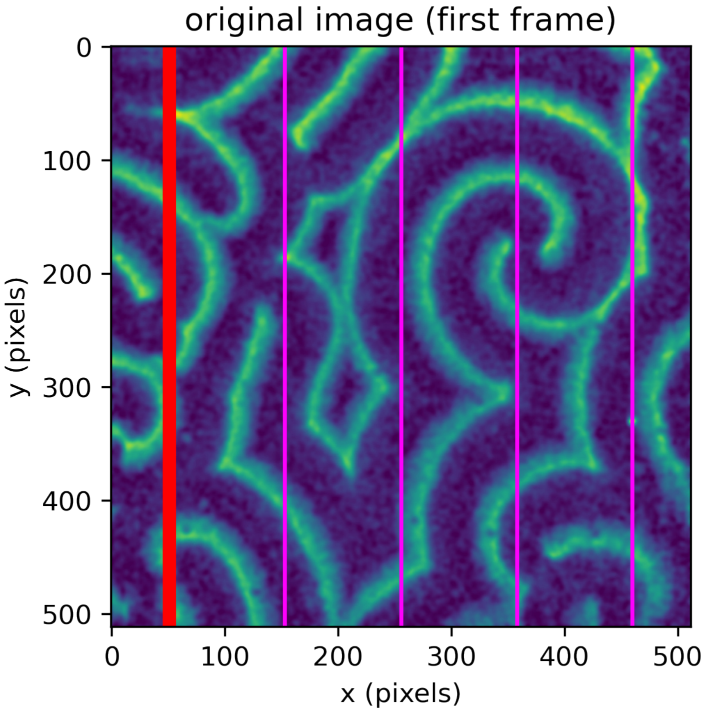


```
[14]: selection_reslice_y = f"{selection}_y_resliced"
(
    crmx_storage_y,
    slices2analyze_y,
    fig,
    ax_corr,
    ax_orig,
) = correlation_tools.get_temporal_correlation_matrixes(
    MinDE_shift_ty,
    "y",
    kymoband,
    reps_per_kymostack,
    demo=True,
)
```

```
Analyzing t-y slices for x = [ 51 153 256 358 460]
```

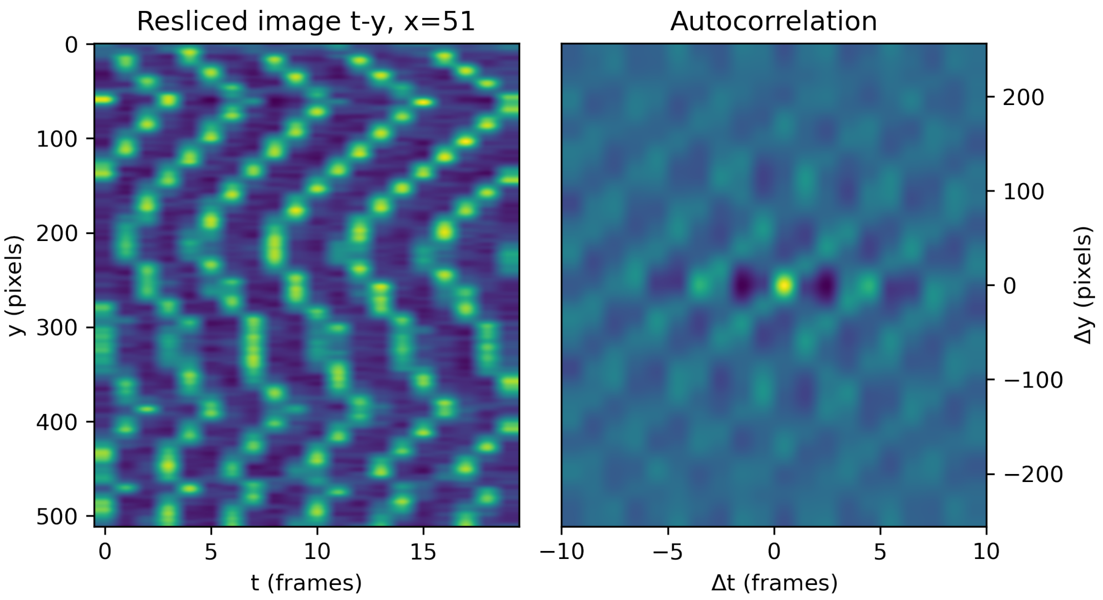


[15]:
```python
(
    first_min_pos_y,
    first_min_val_y,
    first_max_pos_y,
    first_max_val_y,
    fig,
    ax,
) = correlation_tools.analyze_temporal_profiles(
    "y", crmx_storage_y, slices2analyze_y, demo=True
)
```

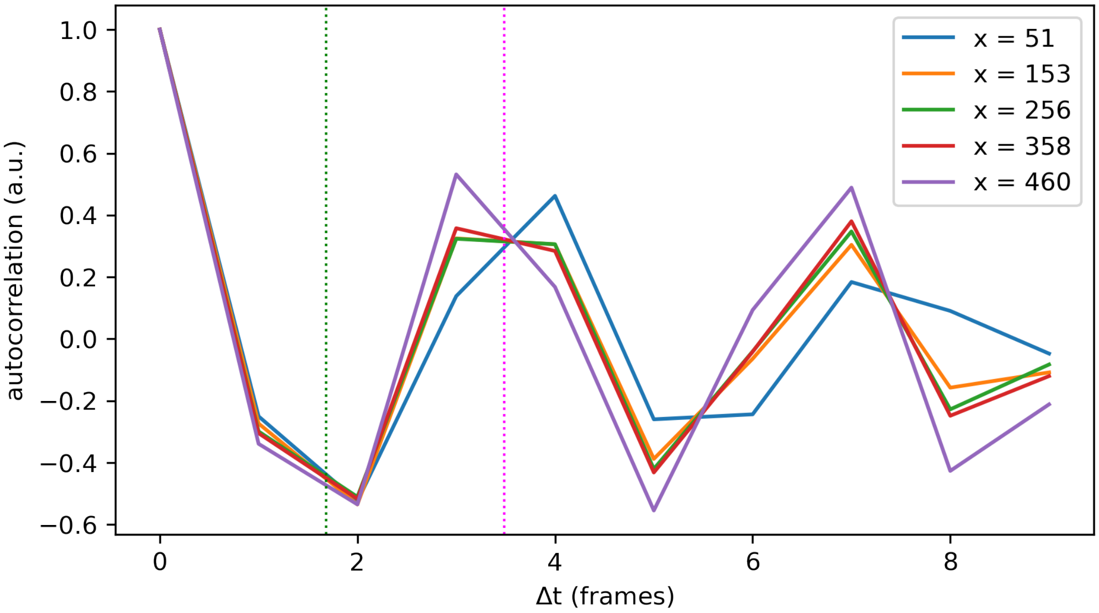


Finally, the parameters collected from all used slices can be averaged to identify the predominant, “global” oscillation period $\tau$ of the pattern:

[16]:
```
# output characteristic parameters
print(
    f"Mean position of first valley: {np.mean(np.append(first_min_pos_x,␣
 ↪first_min_pos_y)):.02f}"
)
print(
    f"Mean position of first peak: {np.mean(np.append(first_max_pos_x,␣
 ↪first_max_pos_y)):.02f} (oscillation period in frames)"
)
```

```
Mean position of first valley: 1.67
Mean position of first peak: 3.49 (oscillation period in frames)
```

### DEMO_MinDE_local_analysis

June 24, 2022

#### 1 Wave crest propagation speed of MinDE patterns

*Jacob Kerssemakers, Sabrina Meindlhumer, Cees Dekker lab, 2022*

The flowing, semi-periodic patterns of MinDE pose some challenges to quantification. Here, we illustrate a systematic approach to determine wave crest propagation velocities $\vec{v} = (v_x,\ v_y)$ for individual wave crest points. Our approach includes the identification of wave crests (using optical flow image analysis tools) and frame-wise comparison of crest positions to determine their local propagation speed. Unlike global analysis (compare our illustration for extraction of global parameters using autocorrelation methods), this approach has the potential to directly yield large distributions of parameters, associated with their $(x,\ y)$ coordinates within the field-of-view.

**READ-ME:** Cells in this notebook need to be executed sequentially. Upon starting to explore this notebook, click the double-arrow symbol above (*Restart the kernel, then re-run the whole notebook*) and hit "Restart" to ensure all required packages are loaded. After that, the notebook will take a few moments to set up, and figures/plots will re-appear one by one. At distinct positions in the notebook, the user is invited to change numeric input. After doing so, the notebook needs to be executed anew at least from this point on for changes to be applied. The notebook can be re-run from any given point onwards by clicking *Run* in the menu-bar above, and selecting *Run Selected Cell and All Below*. Alternatively, the double-arrow symbol can be used again, which will re-run the notebook from the start. This will take a moment longer, but will have the same effect.

##### 1.1 Setup

Import of standard modules, the used flow field analysis method (Horn-Schunck) and assisting custom-made modules. Needs to be executed at least once to ensure correct functionality (see instructions above).

[1]:
```
from pathlib import Path
import numpy as np
from skimage import io
import matplotlib.pyplot as plt
from cv2 import filter2D

from pyoptflow import HornSchunck
from min_analysis_tools.get_auto_halfspan import get_auto_halfspan
from min_analysis_tools import get_data, min_de_patterns_crests, peak_profile
from min_analysis_tools.local_velocity_analysis import local_velocity_analysis
```

```
# Reload modules automatically before executing code
%reload_ext autoreload
%autoreload 2

# Figure quality
import matplotlib as mpl
mpl.rcParams['figure.dpi'] = 300
```

##### 1.2 Select example

Choose an example from the provided stacks in the list below: (1) Simulated spiral (2) Min spiral (example data) (3) Min southeast-directed traveling waves (example data) (4) Min west-directed traveling waves (example data) (5) Min large stitched pattern (example data) (6) Min horizontally stitched pattern (example data) Choose the example by setting the variable *selection* in the code-box below. The notebook needs to be re-run (at least from this point onwards) for changes to be applied.

[2]:
```
selection = 2  # set to 1 ... 6
```

[3]:
```
if selection not in np.arange(1, 7):
    print("Invalid selection. Set to selection 1 (Simulated spiral).")
    selection = 1
    MinDE_st_original = get_data.generate_pattern(
        lambda_t=20, lambda_x=1, size=512, N_frames=50, demo=False
    )
    zm_lo = 45
    zm_hi = 70
elif selection == 1:  # "Simulated spiral"
    MinDE_st_original = get_data.generate_pattern(
        lambda_t=20, lambda_x=1, size=512, N_frames=50, demo=False
    )
    zm_lo = 45
    zm_hi = 70
elif selection > 1:
    if selection == 2:  # "Min spiral (example data)"
        stack_path = Path().cwd() / "example_data" / "demo_spiral.tif"
        zm_lo = 75
        zm_hi = 100
    if selection == 3:  # "Min southeast-directed traveling waves (example data)"
        stack_path = Path().cwd() / "example_data" / "demo_southeast.tif"
        zm_lo = 75
        zm_hi = 100
    if selection == 4:  # "Min west-directed traveling waves (example data)"
        stack_path = Path().cwd() / "example_data" / "paper_west_E.tif"
        zm_lo = 150
        zm_hi = 175
    if selection == 5:  # "Min large stitched pattern (example data)"
```

```
        stack_path = Path().cwd() / "example_data" / "demo_square_stitch.tif"
        zm_lo = 75
        zm_hi = 100
    if selection == 6:  # "Min horizontally stitched pattern (example data)"
        stack_path = Path().cwd() / "example_data" / "demo_horizontal_stitch.tif"
        zm_lo = 65
        zm_hi = 90
    MinDE_st_original = io.imread(stack_path)
nt, ny, nx = np.shape(MinDE_st_original)
auto_halfspan = get_auto_halfspan(
    MinDE_st_original, frames_to_analyse=10, verbose=False
)

fig, ax = plt.subplots()
ax.imshow(MinDE_st_original[0, :, :])
ax.set_xlabel("x (pixels)")
ax.set_ylabel("y (pixels)")
plt.show()

print(f"Current selection: {selection}")
```

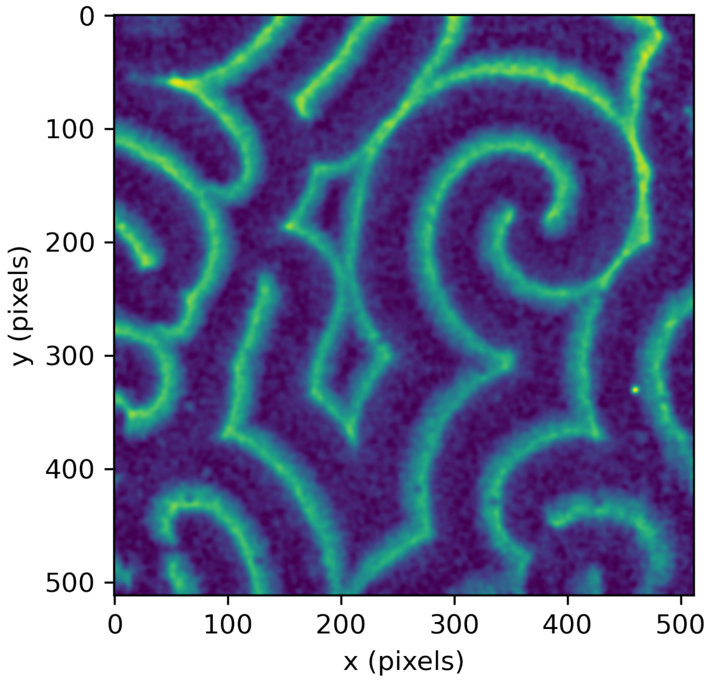


```
Current selection: 2
```

Note that depending on what environment we use, the orientation of $x$- or $y$-components does not always match what you see by eye when opening the image stack with an image viewer, such as Fiji. When relying on the indexing used by NumPy and representation by Matplotlib, we can ensure that the final vector direction matches the apparent directionality by performing some simple transformations on the stack before starting.

[4]:
```
MinDE_st = min_de_patterns_crests.adjust_stack_orientation(MinDE_st_original)
```

[5]:
```
fig, ax = plt.subplots()
ax.imshow(MinDE_st[0, :, :])
ax.set_xlabel("x (pixels)")
ax.set_ylabel("y (pixels)")
plt.show()
```

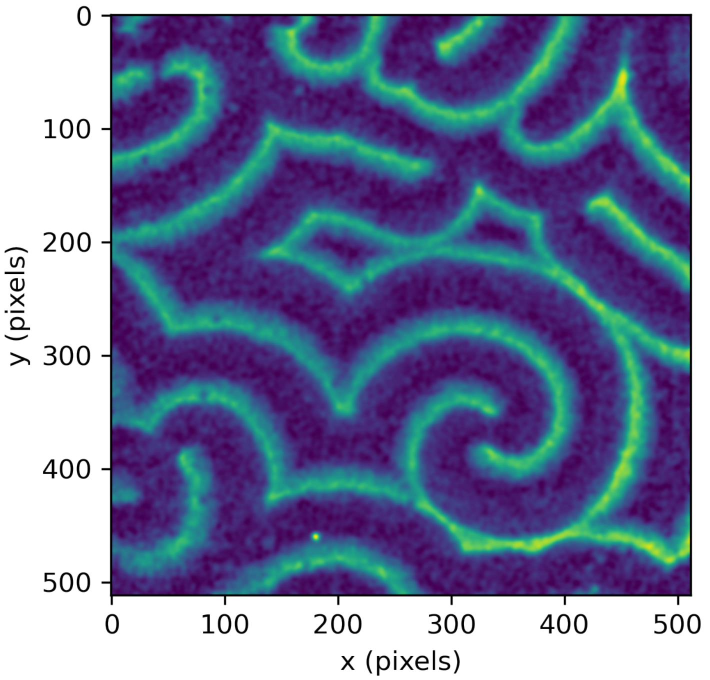


An important parameter to define is *halfspan*, the meaning of which will be explained further below. As a rule of thumb, *halfspan* should be set to approximately half the global wavelength. This can be determined from global analysis or estimated from visual inspection of the pattern. For our demo, we suggest default values for *halfspan*, determined via spatial autocorrelation analysis in the code-box below the example selection. To try other values for *halfspan*, replace "auto_halfspan" by a numeric value (length in pixel). Choosing a value that strongly deviates from the recommended one can lead to erroneous results. Another parameter is *sampling_width*, will later on define the distance between sampling points when comparing crest point position between frames. These parameters can be set in the code-box below. The notebook needs to be re-run (at least from this point on) for changes to apply.

[6]:
```
halfspan = auto_halfspan  # default: auto_halfspan
sampling_width = 1  # default: 1
```

[7]:
```
print(f"Current halfspan: {halfspan} pixels")
print(f"Current sampling_width: {sampling_width} pixels")
```

```
Current halfspan: 41 pixels
Current sampling_width: 1 pixels
```

In the following, we work through the given image stack by performing optical-flow analysis on pairs of sequential images and using this information to identify wave crests.

##### 1.3 Flow field analysis

We build two smoothed image stacks, first one with a finer smoothing kernel, then one with a rather large kernel. The strongly-smoothed stack (*_smz_flow) *is used only for flow analysis (Horn-Schunck), the lightly-smoothed stack (*_smz*) will later on be used for peak detection. The value for light and strong smoothing can be set in the code-box below. Changing these values will lead to results of different quality, and optimal values may differ from file to file. The notebook needs to be re-run (at least from this point on) for change to apply.

[8]:
```
kernel_size_general = 15  # for general (light) smoothing
kernel_size_flow = 35  # stronger smoothing for optical flow analysis
```

[9]:
```
# build kernel for first smoothing step (for processing)
general_kernel = np.ones((kernel_size_general, kernel_size_general), np.float32)␣
 ↪/ (
    kernel_size_general**2
)

# build kernel for obtaining flow pattern
flow_kernel = np.ones((kernel_size_flow, kernel_size_flow), np.float32) / (
    kernel_size_flow**2
)

# create lightly and strongly smoothed images
im0_raw = MinDE_st[0, :, :]
im1_raw = MinDE_st[1, :, :]
im0_smz = filter2D(im0_raw, -1, general_kernel)
im1_smz = filter2D(im1_raw, -1, general_kernel)
im0_smz_flow = filter2D(im0_smz, -1, flow_kernel)
im1_smz_flow = filter2D(im1_smz, -1, flow_kernel)

fig, (ax1, ax2, ax3) = plt.subplots(1, 3)
ax1.imshow(im0_raw)
ax1.set_title("Original")
ax1.set_xlabel("x (pixels)")
ax1.set_ylabel("y (pixels)")
ax2.imshow(im0_smz)
ax2.set_title("Lightly smoothed")
ax2.set_xlabel("x (pixels)")
```

```
ax2.set_ylabel("y (pixels)")
ax3.imshow(im0_smz_flow)
ax3.set_title("Strongly smoothed")
ax3.set_xlabel("x (pixels)")
ax3.set_ylabel("y (pixels)")
fig.tight_layout()
plt.show()
```

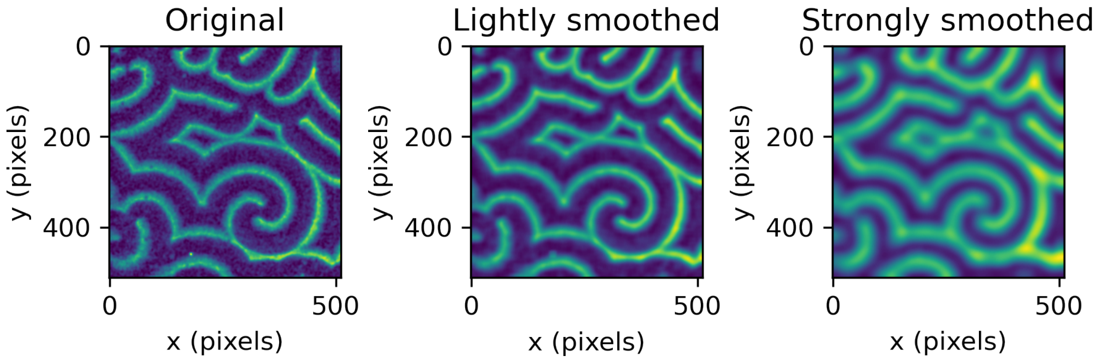


For each sequential pair of images in our stack, we first perform flow analysis using Horn-Schunck algorithm, which gives us an estimate of the local mass flow of fluorescent intensity between subsequent image frames We obtain this information in the form of flow field vector maps describing the local flow components in $x$- and $y$-direction. Note that this is still local flow and not the actual wave propagation. In the map below, the magnitude of each position's corresponding flow vector is shown as image brightness.

[10]:
```
flow_x, flow_y = HornSchunck(im0_smz_flow, im1_smz_flow, alpha=100, Niter=100)
```

[11]:
```
# Show the optical flow magnitude
flow_mag = np.hypot(flow_x, flow_y)
fig, ax = plt.subplots()
ax.imshow(flow_mag)
ax.set_title("pattern flow magnitude")
ax.set_xlabel("x (pixels)")
ax.set_ylabel("y (pixels)")
plt.show()
```

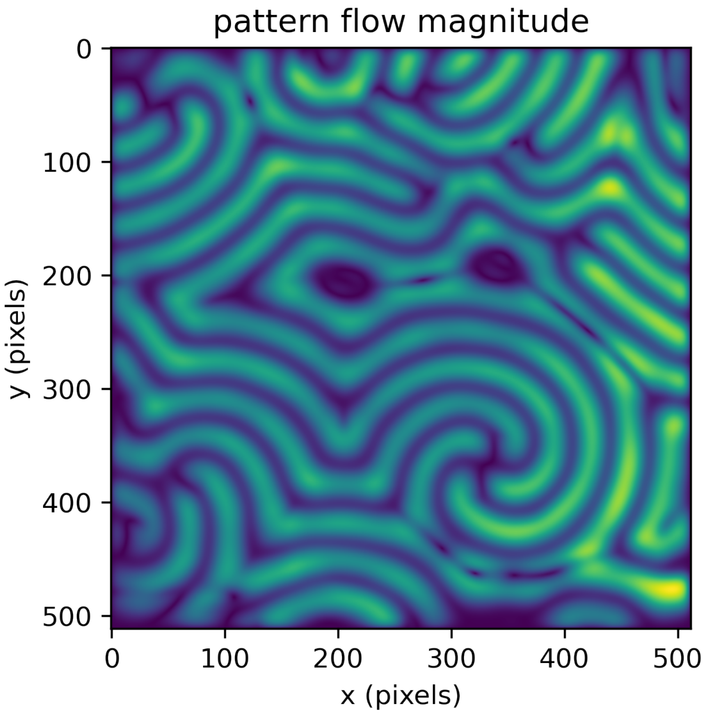


##### 1.4 Crest detection

Next, we can use this optical flow analysis results to identify wave crests. We do this by starting from a given pixel position, using the corresponding vector obtained from optical flow analysis and comparing the pixel's intensity to the one at a position located a short distance along the direction of the flow vector. That allows us to divide the image in areas where fluorescent intensity is decreasing in the direction of flow (corresponding to the "front" of a wave) or increasing (its "wake"). This binary division can be represented as a wavesign-image, representing the front regions as bright and wake regions as dark areas.

[12]:
```
wavesign_im = min_de_patterns_crests.get_rise_or_fall(flow_x, flow_y, im0_smz)
```

[13]:
```
fig, ax = plt.subplots()
ax.imshow(wavesign_im)
ax.set_xlabel("x (pixels)")
ax.set_ylabel("y (pixels)")
plt.show()
```

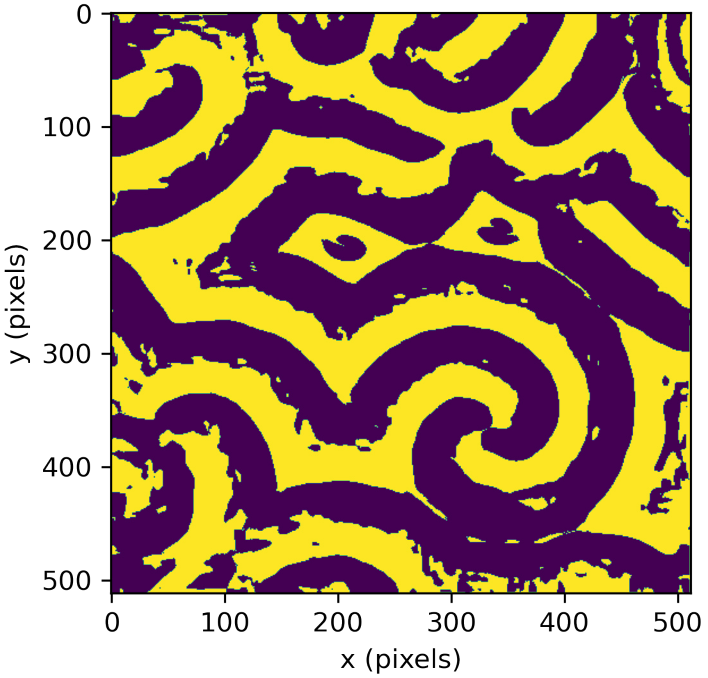


Using this image, we obtain the crests, defined as the points that lie between front or wake areas *and* where the intensity is above average (since otherwise, we would also get the "valleys"). Each crest point comes with $(x, y)$ coordinates and a unit vector $(w_x, w_y)$ perpendicular to the local crest line, in the direction of wave propagation.

[14]:
```
(
    crests_x,
    crests_y,
    forward_wavevector_x,  # w_x
    forward_wavevector_y,  # w_y
) = min_de_patterns_crests.get_crests(wavesign_im, im0_smz, halfspan / 2)
```

In the representation below, the identified wave crests are shown in red, and their direction of propagation is indicated in blue. Crest points close the edges of the frame are discarded to avoid edge-effects.

[15]:
```
fig, ax = plt.subplots()
span = 10
cx2 = crests_x + span * forward_wavevector_x
cy2 = crests_y + span * forward_wavevector_y
ax.imshow(im0_raw)
ax.plot(crests_x, crests_y, "ro", markersize=3)
ax.plot(cx2, cy2, "bo", markersize=1)
ax.set_xlabel("x (pixels)")
ax.set_ylabel("y (pixels)")
```

```
plt.show()
```

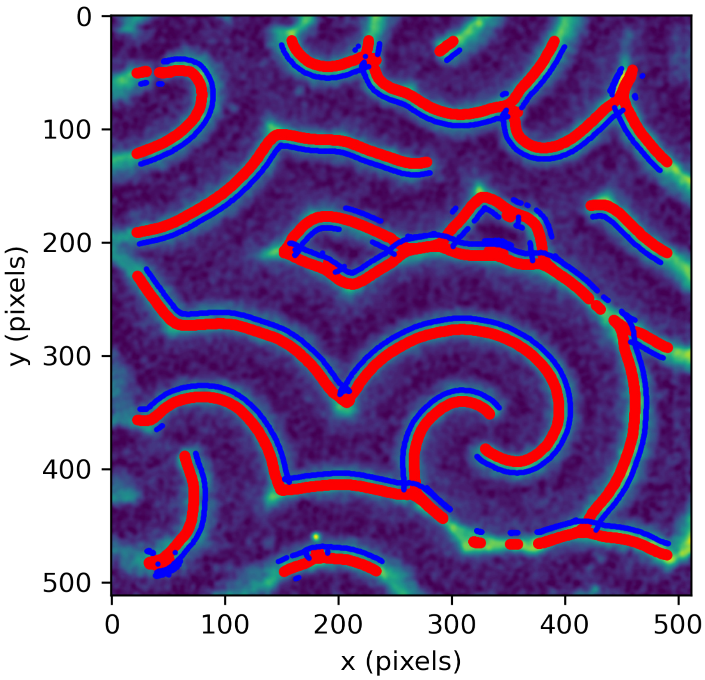


#### 1.5 Profile mapping

We can use the wave vectors at the identified crest positions to estimate the local propagation of the pattern. For illustration, we plot a corner of the pattern. Again, the detected crest points are shown in red, an indication of local propagation direction in blue.

[16]:
```
fig, ax = plt.subplots()
zm = 200
Le = len(crests_x)
span = 10
cx1 = crests_x
cy1 = crests_y
cx2 = cx1 + span * forward_wavevector_x
cy2 = cy1 + span * forward_wavevector_y
zm_ix = np.argwhere((cx1 < zm) & (cy1 < zm))
ax.imshow(im0_raw[0 : zm - 1, 0 : zm - 1])
ax.plot(cx1[zm_ix], cy1[zm_ix], "ro", markersize=3)
ax.plot(cx2[zm_ix], cy2[zm_ix], "bo", markersize=1)
ax.set_xlabel("x (pixels)")
ax.set_ylabel("y (pixels)")
plt.show()
```

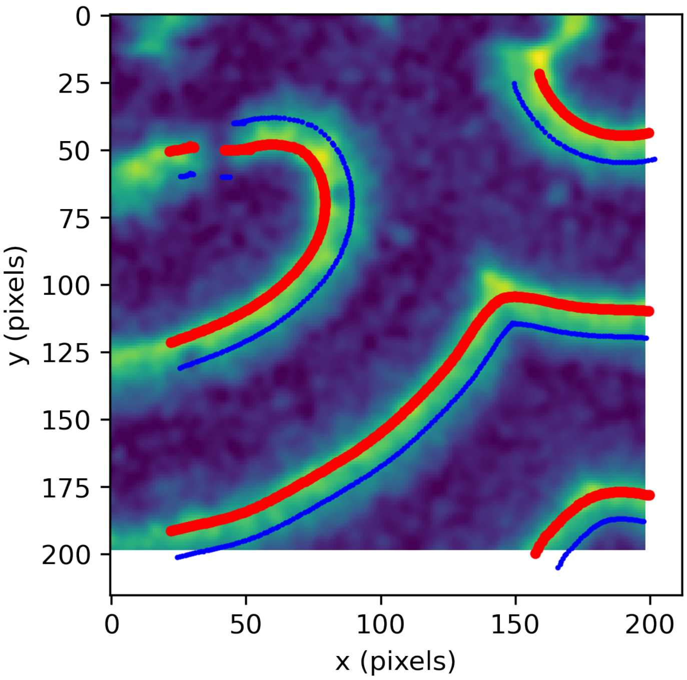


For each crest point and its accompanying wave vector, we can now obtain a cross-section profile by two-dimensional interpolation. For efficiency, this is done as a grid-style 2D interpolation for all the crest points together. Left panel: For clarity, we show a zoomed-in section of the pattern and the overlaying crests (red) and cross-section samplings (black). Right panel: The corresponding sampling map.

[17]:
```
(profile_map1, xxgrid, yygrid) = min_de_patterns_crests.sample_crests(
    im0_smz,
    crests_x,
    crests_y,
    forward_wavevector_x,
    forward_wavevector_y,
    halfspan,
    sampling_width,
)
```

[18]:
```
# Show a zoom-in of the result, specified by the variables zm_lo and zm_hi
# zm_lo and zm_hi are defined in the code-box below the selection above

rr, cc = np.shape(im0_raw)
zm_ix1 = np.argwhere(
    (crests_x > zm_lo) & (crests_y > zm_lo) & (crests_x < zm_hi) & (crests_y <␣
 ↪zm_hi)
)
zm_ix2 = np.argwhere(
```

```
    (xxgrid.ravel() > zm_lo)
    & (yygrid.ravel() > zm_lo)
    & (xxgrid.ravel() < zm_hi)
    & (yygrid.ravel() < zm_hi)
)
fig, (ax_1, ax_2) = plt.subplots(1, 2)

ax_1.imshow(im0_raw[zm_lo:zm_hi, zm_lo:zm_hi])
ax_1.plot(
    xxgrid.ravel()[zm_ix2] - zm_lo,
    yygrid.ravel()[zm_ix2] - zm_lo,
    "ko",
    markersize=1,
)
ax_1.plot(
    crests_x[zm_ix1] - zm_lo,
    crests_y[zm_ix1] - zm_lo,
    "ro",
    markersize=5,
)
ax_1.set_xlabel("x (pixels)")
ax_1.set_ylabel("y (pixels)")

ax_2.imshow(
    profile_map1.T[:, zm_ix1],
    aspect="auto",
    origin="lower",
    extent=[0, crests_x[zm_ix1].size, -halfspan, halfspan],
)
ax_2.set_xlabel("crest index (within zoom-in)")
ax_2.set_ylabel("position (pixels)")

fig.tight_layout()
plt.show()
```

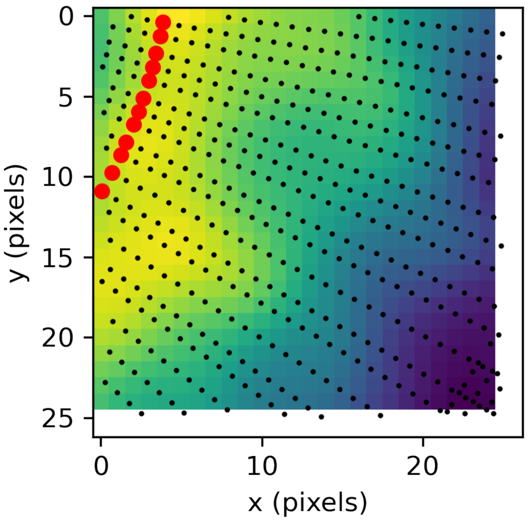


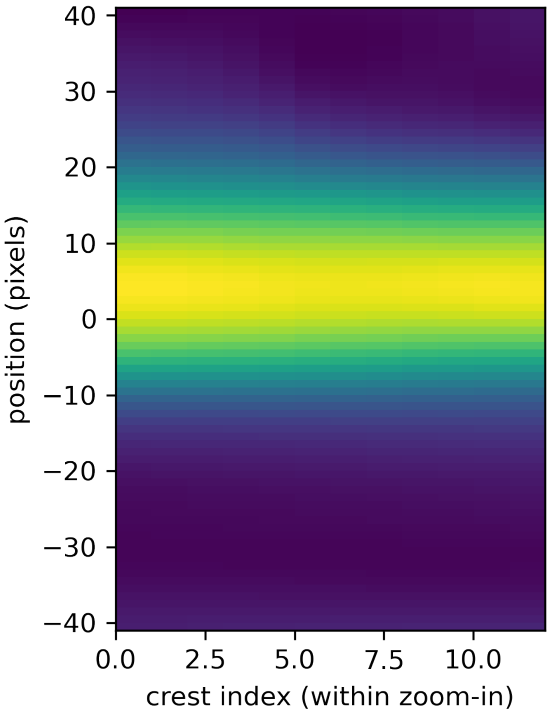


The same sampling grid is now used for the next image frame. As can be seen in the second sampling map, the (upwards) shift of all profile maxima reflects the motion of the crests from one frame to the next. Note that the parameter *halfspan* defines the distance (plus/minus) starting from the crest position of the first frame, within which intensities are sampled (at density *sampling_ width*). If there is no shifted intensity profile visible in the sampling map below, it could be that a higher value for *halfspan* has to be chosen.

[19]:
```
(profile_map2, xxgrid, yygrid) = min_de_patterns_crests.sample_crests(
    im1_smz,
    crests_x,
    crests_y,
    forward_wavevector_x,
    forward_wavevector_y,
    halfspan,
    sampling_width,
)
```

[20]:
```
# Show a zoom-in of the result, specified by the variables zm_lo and zm_hi
# zm_lo and zm_hi are defined in the code-box below the selection above

rr, cc = np.shape(im1_raw)
zm_ix1 = np.argwhere(
    (crests_x > zm_lo) & (crests_y > zm_lo) & (crests_x < zm_hi) & (crests_y <␣
↪zm_hi)
```

```
)
zm_ix2 = np.argwhere(
    (xxgrid.ravel() > zm_lo)
    & (yygrid.ravel() > zm_lo)
    & (xxgrid.ravel() < zm_hi)
    & (yygrid.ravel() < zm_hi)
)
fig, (ax_1, ax_2) = plt.subplots(1, 2)

ax_1.imshow(im1_raw[zm_lo:zm_hi, zm_lo:zm_hi])
ax_1.plot(
    xxgrid.ravel()[zm_ix2] - zm_lo,
    yygrid.ravel()[zm_ix2] - zm_lo,
    "ko",
    markersize=1,
)
ax_1.plot(
    crests_x[zm_ix1] - zm_lo,
    crests_y[zm_ix1] - zm_lo,
    "ro",
    markersize=5,
)
ax_1.set_xlabel("x (pixels)")
ax_1.set_ylabel("y (pixels)")

ax_2.imshow(
    profile_map2.T[:, zm_ix1],
    aspect="auto",
    origin="lower",
    extent=[0, crests_x[zm_ix1].size, -halfspan, halfspan],
)
ax_2.set_xlabel("crest index (within zoom-in)")
ax_2.set_ylabel("position (pixel)")

fig.tight_layout()
plt.show()
```

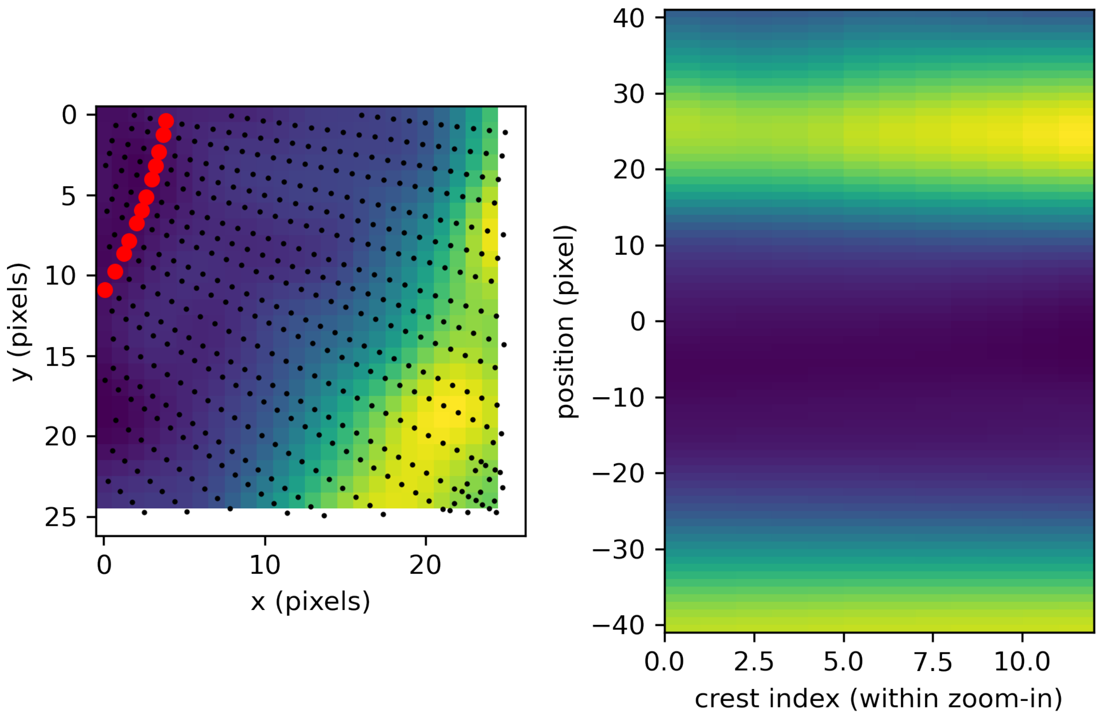


##### 1.6 Velocity tracking

The two maps offer two profiles per crest position, one for the current frame (frame 1) and one for the next (frame 2). For each profile, a sub-unit maximum position is determined by a multi-point parabolic fit around the maximum. Since the cross-section was taken perpendicular to the wave crest, the shift between the two maxima reflects the shift of the crest peak position perpendicular to the local wave orientation. Thus, this shift equals a crest velocity in pixels per frame. The translocation distance $d$ (in distance units) from peak to peak divided the time $\Delta t$ that passed between the acquisition (in time units) on sequential frames then yields the velocity magnitude: $v = \frac{d}{\Delta t}$ Combined with the direction of the normalized flow vectors $(w_x, w_y)$, this yields a vector $\vec{v}$ for each individual crest position $(x, y)$. The parameter $look\_ahead = 1$ defines that we are considering peaks located in front of the current one, that is, in propagation direction (as determined from optical flow analysis). If no local maximum can be determined in the direction of interest, the crest point is discarded. In other applications of the local analysis pipeline (like DE distance detection, when looking for the position of a MinE wave running behind a MinD wave), it can make sense to look towards the other direction.

[21]:
```
velocities = min_de_patterns_crests.compare_crestmaps(
    profile_map1, profile_map2, sampling_width, look_ahead=1
)
```

[22]:
```
print(f"Determined velocities for {np.size(velocities)} crest points.")
bad_count = 0
bad_indices = []
```

```
for ind, vel in enumerate(velocities):
    if not vel >= 0:
        bad_indices.append(int(ind))
        bad_count = bad_count + 1
velocities = np.delete(velocities, bad_indices, 0)
profile_map1 = np.delete(profile_map1, bad_indices, 0)
profile_map2 = np.delete(profile_map2, bad_indices, 0)
xxgrid = np.delete(xxgrid, bad_indices, 0)
yygrid = np.delete(yygrid, bad_indices, 0)
print(f"Discarded {bad_count} peaks due to failed peak detection.")
```

```
Determined velocities for 3441 crest points.
Discarded 390 peaks due to failed peak detection.
```

Below, the profile along the highlighted line will be shown in red for frame 1 and in blue for frame 2. The maximum identified will be shown as black dot. The crest index to be shown can be set below (notebook needs to be re-run from this point on for changes to apply).

[23]:
```
length = np.shape(profile_map1)[0]
print(
    f"Considered wave crest points: {length} (set crest index to integer number␣
 ↪from 0 to {length-1})"
)
```

```
Considered wave crest points: 3051 (set crest index to integer number from 0 to
3050)
```

[24]:
```
crest_index = 1000  # crest index for viewing, can be changed here
```

[25]:
```
# show position of profile to be displayed on the image
fig, (ax1, ax2) = plt.subplots(1, 2)
ax1.imshow(im0_raw)
ax1.plot(xxgrid[crest_index], yygrid[crest_index], color="white", linewidth=3)
ax1.set_title("frame 1 (red trace)")
ax1.set_xlabel("x (pixels)")
ax1.set_ylabel("y (pixels)")
ax2.set_title("frame 2 (blue trace)")
ax2.imshow(im1_raw)
ax2.plot(xxgrid[crest_index], yygrid[crest_index], color="white", linewidth=3)
ax2.set_xlabel("x (pixels)")
ax2.set_ylabel("y (pixels)")
fig.tight_layout()

# identify peak positions for this particular crest location
prf1 = profile_map1[crest_index, :]
prf2 = profile_map2[crest_index, :]
peak_shift, x1, y1, x2, y2 = min_de_patterns_crests.compare_profiles(
    prf1, prf2, sampling_width, look_ahead=1, return_peaks=True
```

```
)
print(len(prf1))

# create shifted axis to have first peak close to center and plot
x_axis_shift = np.arange(-halfspan, halfspan, sampling_width)

# plot example profile
fig, ax = plt.subplots()
ax.set_title(f"Profile plot for crest index {crest_index}")
ax.plot(x_axis_shift, prf1, "ro-", markersize=2)
ax.plot(x_axis_shift, prf2, "bo-", markersize=2)
ax.plot(x1 * sampling_width - halfspan, y1, "ko", markersize=6)
ax.plot(x2 * sampling_width - halfspan, y2, "ko", markersize=6)
ax.set_xlabel("distance (pixels)")
ax.set_ylabel("intensity (a.u.)")
ax.set_xlim([-halfspan, halfspan - sampling_width])
fig.tight_layout()
plt.show()
```

82

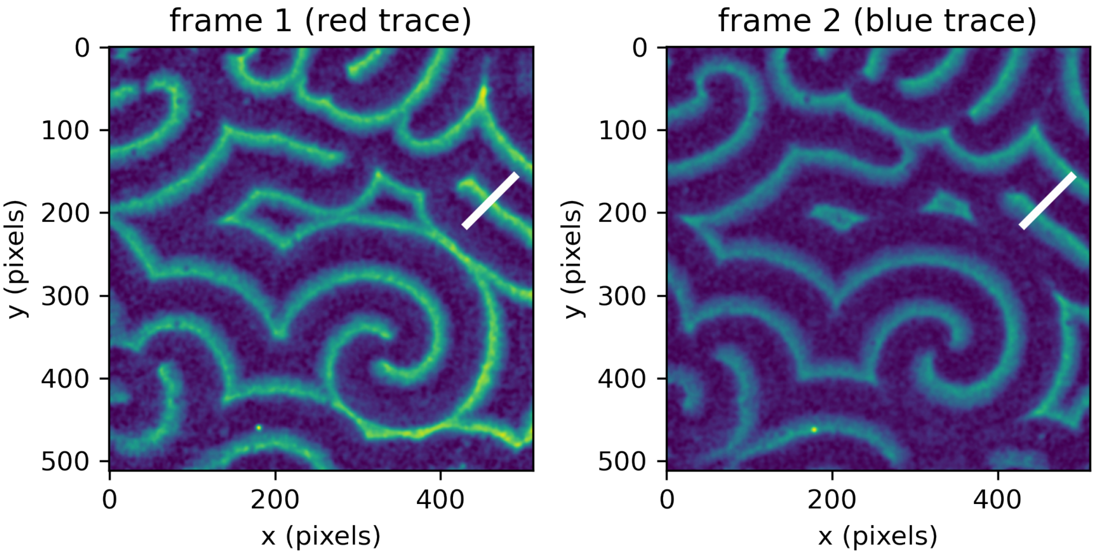

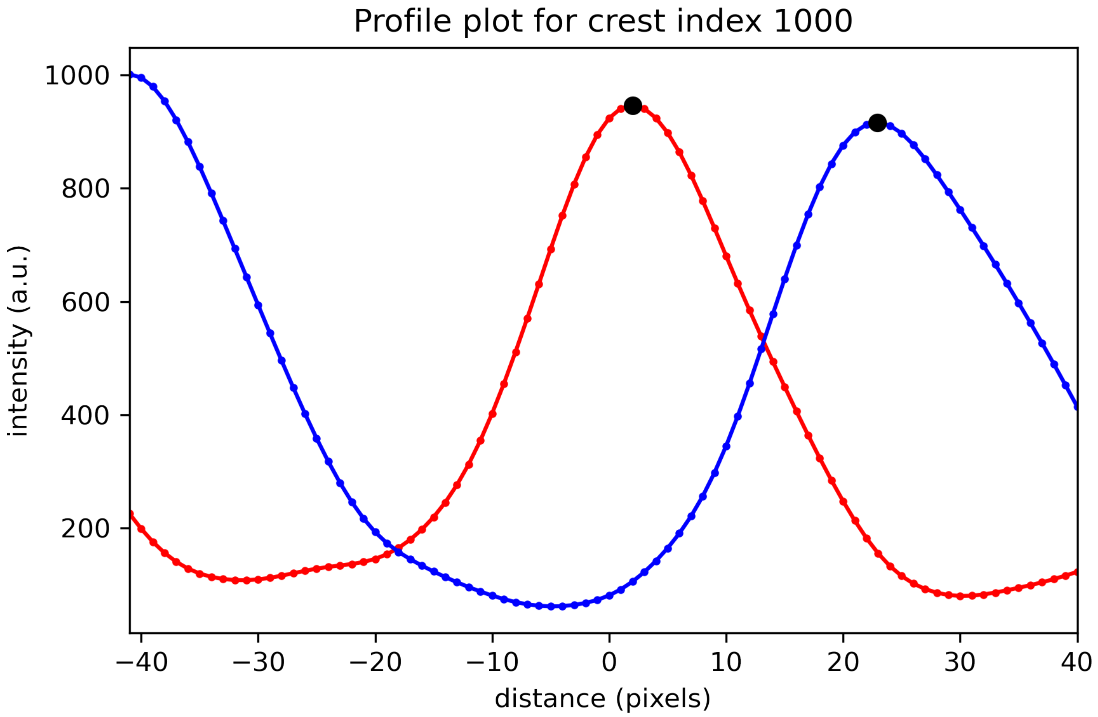


##### 1.7 Velocity distribution

The analysis routine presented so far can be performed for all pairs of frames within a given image stack. Collecting these results yields a distribution of velocity vectors, which can be represented in different ways, depending on what information (magnitude, directionality) is of interest.

[26]:
```
(
    velocities,  # velocity magnitude
    forward_wavevector_x,  # unit vector w_x
    forward_wavevector_y,  # unit vector w_y
    all_wheels,  # data generated for velocity wheel (2D histogram)
    crests_x,
    crests_y,
    framenr,
    max_x1,
    max_y1,
    max_x2,
    max_y2,
    fig,
    ax_wheel,
    ax_sum,
) = local_velocity_analysis(
    MinDE_st_original,
    frames_to_analyse=5,  # use first ... frames
```

```
    halfspan=halfspan,
    sampling_width=sampling_width,
    edge=40,  # width of velocity wheel (2D histogram) and max of magnitude␣
↪histogram
    bins_wheel=40,  # number of bins (horizontal/vertical) for velocity wheel␣
↪(2D histogram)
    binwidth_sum=1,  # binwidth for velocity mangitude histogram
    kernel_size_general=kernel_size_general,  # kernel for first smoothing step
    kernel_size_flow=kernel_size_flow,  # kernel for additional smoothing step
    look_ahead=1,  # 1 -> look in propagation direction, -1 -> against it
    demo=True,  # True -> return figure handles
)
plt.show()
```

```
Analysing 5 frames
Working frame 0 to 1
Working frame 1 to 2
Working frame 2 to 3
Working frame 3 to 4
peak velocity magnitude: 19.00 pixels/frame
FWHM velocity magnitude: 5.00 pixels/frame
Median velocity magnitude: 19.90 pixels/frame
```

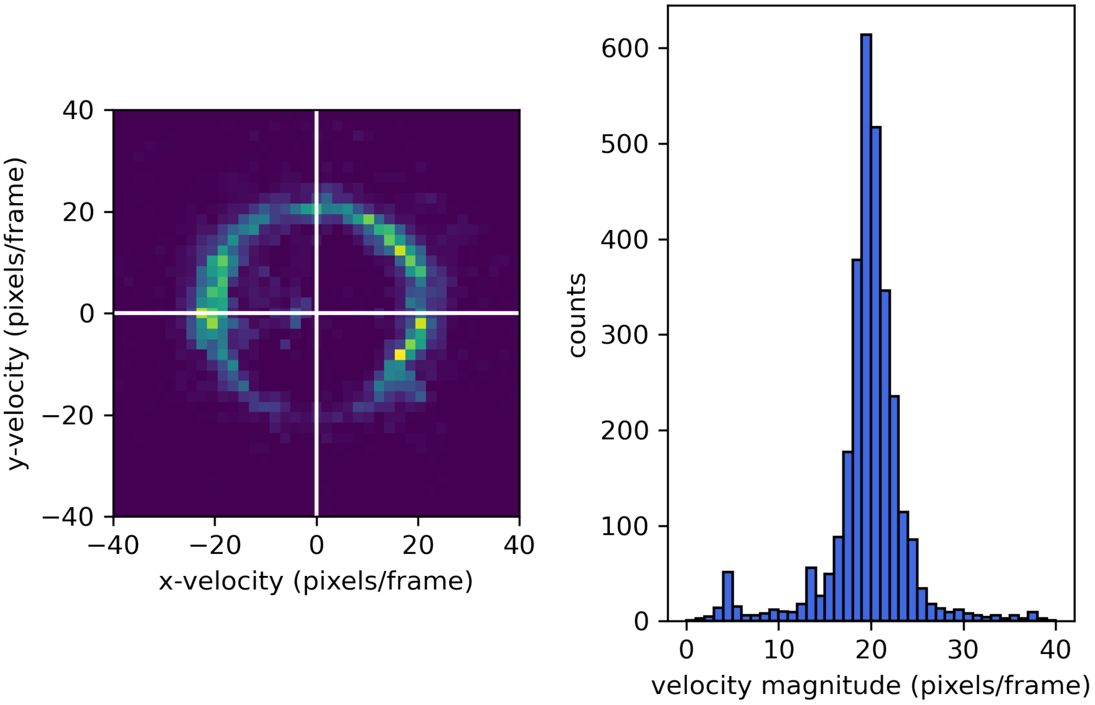
